# Decoding Salience: A Functional Magnetic Resonance Imaging Investigation of Reward and Contextual Unexpectedness in Memory Encoding and Retrieval

**DOI:** 10.1101/2024.05.27.596071

**Authors:** Yeo-Jin Yi, Michael C. Kreißl, Oliver Speck, Emrah Düzel, Dorothea Hämmerer

## Abstract

The present study investigated the neuromodulatory substrates of salience processing and its impact on memory encoding and behaviour, with a specific focus on two distinct types of salience: reward and contextual unexpectedness. 46 participants performed a novel task paradigm modulating these two aspects independently and allowing for investigating their distinct and interactive effects on memory encoding while undergoing high resolution fMRI. By using advanced image processing techniques tailored to examine midbrain and brainstem nuclei with high precision, our study additionally aimed to elucidate differential activation patterns in subcortical nuclei in response to reward-associated and contextually unexpected stimuli, including distinct pathways involving in particular dopaminergic modulation. We observed a differential involvement of the ventral striatum, substantia nigra and caudate nucleus, as well as a functional specialisation within the subregions of the cingulate cortex for the two salience types. Moreover, distinct subregions within the substantia nigra in processing salience could be identified. Dorsal areas preferentially processed salience related to stimulus processing (of both reward and contextual unexpectedness) versus ventral areas were involved in salience-related memory encoding (for contextual unexpectedness only). These functional specialisations within SN are in line with different projection patterns of dorsal and ventral SN to brain areas supporting attention and memory, respectively. By disentangling stimulus processing and memory encoding related to two salience types, we hope to further consolidate our understanding of neuromodulatory structures’ differential as well as interactive roles in modulating behavioural responses to salient events.

## 1 Introduction

Neuromodulation influences physiological and cognitive functions including memory, attention, and emotion regulation (1–5). Key systems involve the dopaminergic system (substantia nigra [SN] and ventral tegmental area [VTA]; (4,6)), noradrenergic system (locus coeruleus [LC]; (4)), and serotonergic system (raphe nuclei; (7)). Despite their small volume, the midbrain and brainstem harbour the origins of these systems, projecting to different brain regions and affecting various processes such as attention, working memory, and long-term memory (2,8–16).

From animal and human research, it is known that the midbrain and brainstem neuromodulatory systems, especially those responsive to salient events, play a crucial role in memory consolidation (17–23). For instance, evidence from animal studies indicates that it is predominantly the noradrenergic system, and in particular the noradrenergic locus coeruleus in the brainstem, which modulates attention and arousal, enhancing memory retention for novel and aversive events (1,22). On the other hand, dopamine, and in particular the substantia nigra in the midbrain, promotes reward processing and learning, and supports memory encoding for novel or positive events (16,21,23–26). Despite these seemingly straightforward distinctions, animal studies suggest that the separation between noradrenergic and dopaminergic nuclei in processing different types of salience might not be as distinct as previously thought. For example, the processing of novel stimuli, commonly associated with dopaminergic modulation, seems to activate both the locus coeruleus and the substantia nigra, with the latter showing more sustained activity (22). Such co-activations are plausible given the anatomical connections between noradrenergic and dopaminergic cell groups (2). Finally, although perhaps less relevant for functional MRI studies, it is important to consider that neuromodulatory cell groups often release multiple neurotransmitters; for instance, the noradrenergic locus coeruleus also releases dopamine to the hippocampus. Therefore, while fMRI might indicate the involvement of a typically noradrenergic structure, the underlying cognitive effects could be mediated by dopamine (27,28). Taken together, although the influence of event saliency on human memory formation is well recognized, establishing distinct relationships between neuromodulation and enhanced memory for different types of salience such as reward and unexpectedness or novelty in humans is often complicated due to in part overlapping neural substrates (12,21,22,26,29–34). Moreover, the methodological challenges involved in reliably imaging the small neuromodulatory nuclei of the midbrain and brainstem in humans makes it difficult to disentangle and closely inspect the distinct mechanisms (35).

In this study, we aimed to understand the neuromodulatory underpinnings of different types of salience, namely contextual unexpectedness and reward, and their effects on memory encoding. We conducted a two-session experiment in order to separately manipulate the salience effect on memory related to contextual unexpectedness and reward association in the same sample. To effectively investigate the role of neuromodulatory midbrain and brainstem structures in processing salience and encoding memories for salient events, we applied a newly developed MRI data processing approach, which specifically enhances spatial precision in assessing brainstem and midbrain activations, increasing the reliability and significance of our findings (36).

Our study hypothesises that (1) processing different types of saliences and their memory effects will preferentially rely on distinct neural substrates with reward-associated stimuli relying more on dopaminergic networks and unexpectedness-associated stimuli more on predominantly noradrenaline networks (21). Finally, we expect that (2) episodic memory encoding will be facilitated by both reward- and unexpectedness-associated salience, which will be reflected in the enhanced subsequent memory effects for stimuli linked to salience as well as parallel primary support by dopaminergic and noradrenergic networks, respectively.

## 2 Methods

### 2.1 Participants

Fifty healthy younger adults (22 males, age range: 18−31 years, *M±SD*=23.5±2.4) were recruited via the German Center for Neurodegenerative Diseases (DZNE) participant database. MRI eligibility was initially screened via telephone conversations and email. Exclusion criteria included age, history of neurobiological disorders, and the presence of ferromagnetic implants. Each participant was scanned twice as the study compared the effects of two different salience contexts on memory encoding. Three subjects dropped out after the first session due to scheduling issues, thus resulting in a total 47 participants with two scan sessions, i.e. 94 scans. The handling procedures of two-session MRI data are described in detail in the data analysis section (section 2.2.4.) below. All participants provided written informed consent prior to each session. At the end of each experimental visit, they were compensated either 72 Euros or 32 Euros cash depending on the reward context type of the session.

### 2.2 Task design and procedures

#### 2.2.1 Materials

MATLAB R2015b (Mathworks, Sherborn, MA, USA, 2015) and Cogent toolbox (Cogent Graphics, http://www.vislab.ucl.ac.uk/CogentGraphics.html [Accessed May 2018]) were employed for paradigm creation and execution. To provide a comparable range of stimulus memorability, scene images were sourced from the Large-scale Image Memorability dataset (LaMem, (37)) and manually screened to exclude: (1) memorability values outside the 0.4-0.6 range as per LaMem; (2) emotional elements such as blood or sexual content; (3) distinctive face-like features; (4) legible text; (5) animals. Post-screening, images were categorised into four subgroups (public indoor, private indoor, urban outdoor, natural outdoor) to allow for four separate stimulus categories associated with reward or no reward outcomes across the two sessions. The luminance level of all stimuli were set at 50% as stimulus brightness is known to affect pupil dilations, which were concurrently recorded but are not reported here. Background stimuli (binary chequered-noise stimuli) were also set at 50% luminance (Figure 1).

**Figure 1.**
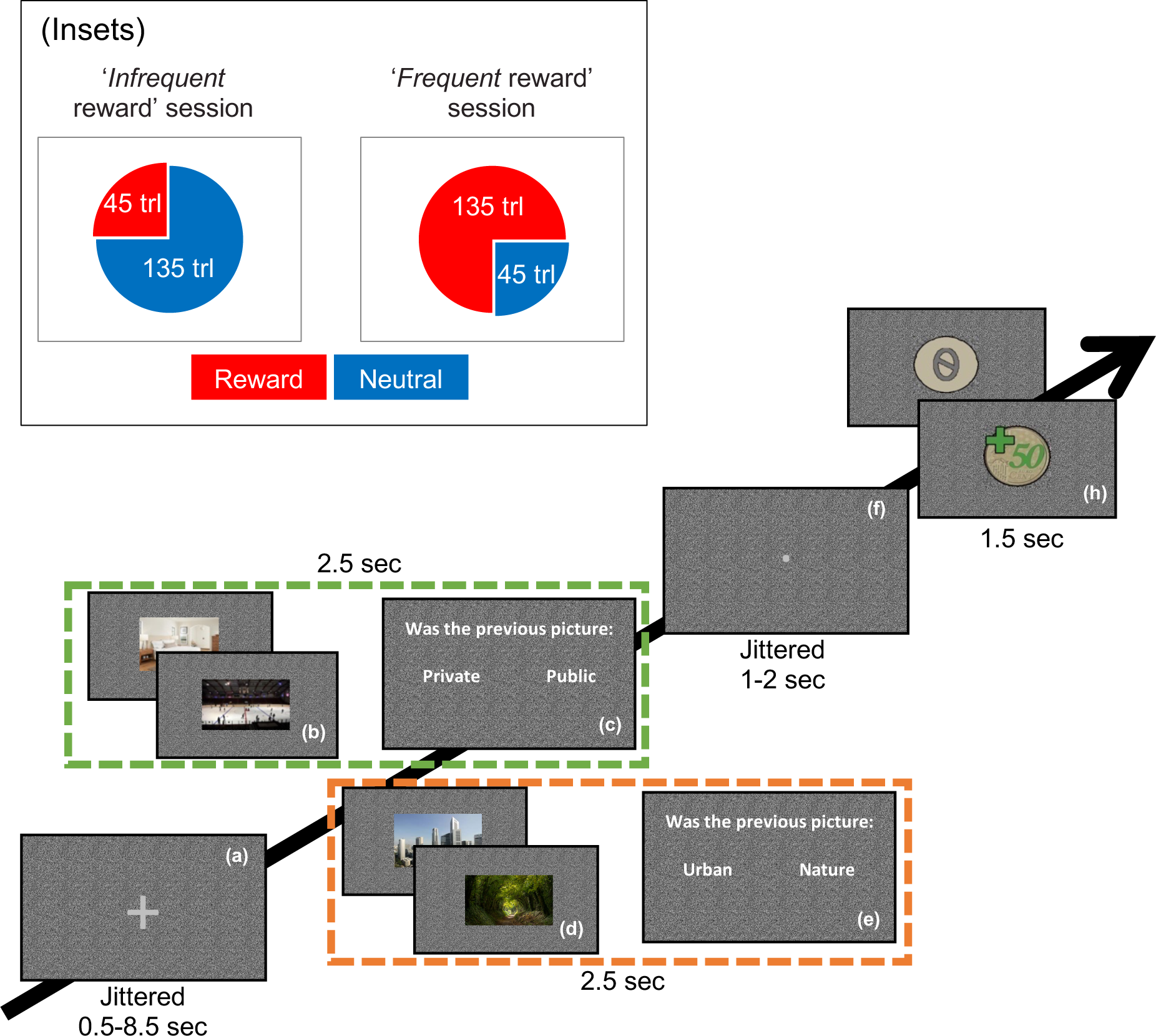
Trial Structure. The figure shows the layout of the stimuli on the screen and the sequence within each trial: (a) baseline, *jittered* between 0.5 and 8.5 seconds in duration; (b, d) scenes to be categorised as either indoor or outdoor, each lasting 2.5 seconds; (c, e) categorisation response, lasting 2 seconds regardless of button input; (f) a subsequent baseline, indicated by a dot, *jittered* between 1 and 2 seconds in duration; (h) 1.5-second feedback presentation, differentiated by the preceding baseline screen. Green and orange dashed boxes indicate example stimulus sets for the two test sessions. Jittered intervals between scene stimuli and feedback were included in order to facilitate investigating functional activations to these two timepoints separately. The **insets** indicate the composition of the infrequent and frequent reward sessions, the order of which was likewise randomised.

#### 2.2.2 Task design and procedures

##### 2.2.2.1 Experimental programme

In our study, we conducted two types of test sessions on separate days within subject to manipulate the reward context, differing in the frequency of reward-associated trials. There were 135 rewarded trials in the ‘frequent reward session’ and 45 in the ‘infrequent reward session,’ with neutral feedback in the remainder (see Figure 1 inset). For example, in one session a subject might encounter an indoor scene stimulus set consisting of private and public scenes, with either private or public scenes randomly assigned as rewarded, while the other category received neutral feedback. In the alternate session (i.e. the second visit), the subject would be presented with an outdoor scene stimulus set, comprised of nature and urban scenes, and either nature or urban scenes would be randomly assigned as rewarded. Across subjects, the order of indoor and outdoor scenes, as well as which category within each set was designated ‘frequent reward’ or ‘infrequent reward’, was randomised. Thus, if private indoor scenes were assigned as ‘frequent reward’ in one session, the rewarded outdoor scene category in the next session would be ‘infrequent’. Subjects were compensated with 50 cents for each rewarded scene. (see also ‘Reward task and memory test’ and Figure 1 below for more details). The interval between the two visits was a minimum of 1 day and maximum of 29 days (*M*=7.33, *SD*=7.56). By manipulating the presentation frequency of rewards in two separate test sessions, the effect of two salience types, reward and contextual unexpectedness, on the following two aspects can be examined: namely a) whether a stimulus is associated with a reward or a neutral outcome, and b) how frequently a stimulus category is presented in the context of a specific session’s reward schedule. In addition, the temporal design of the task was optimized in order to allow for examining functional brain activations to scenes and feedbacks separately. This approach permitted separate assessments of processing salient stimuli as well as the impact of associated feedbacks on memory encoding within the context of different salience types. During each session, functional magnetic resonance imaging (fMRI) as well as structural magnetic resonance imaging (sMRI) was carried out. Pupillometric data were collected simultaneously during fMRI, which will not be reported here.

##### 2.2.2.2 Reward task and memory tests

In the reward task, participants were instructed to sort a picture into two categories per session, one of which was rewarded and one of which was infrequent (Figure 1). All images presented during this encoding task were trial unique. Altogether, in order to distinguish infrequent and frequent as well as rewarded and not rewarded stimuli, four different types of scenes were included across the two sessions: Private or public indoor pictures and urban or nature outdoor pictures (cf. Figure 1). In order to make it easier for participants to differentiate scenes across sessions, one session used indoor scenes, and the other session used outdoor scenes, i.e. indoor and outdoor scenes were never mixed in a session. Within each session, only one scene category (e.g. urban in ‘outdoor session’ or private in ‘indoor session’) was associated with a reward. Reward association of scenes did not change across categories within a session and was deterministic. That is, every incidence of a reward category scene was followed by reward feedback. Which session (‘indoor’ or ‘outdoor’) came first, which scene category was associated with a reward, and of which frequency the reward-associated scenes were presented during the task (‘infrequent’ or ‘frequent reward’ session) were counterbalanced across participants. In this way, no scene category was preferentially associated with a first or second test session or saliency conditions, i.e. frequency or reward, across participants.

Each scan session started with 15-minute sMRI data collection, whole-brain T1, high-resolution T2, and fieldmap. Participants did not perform any tasks during this period and were allowed to close their eyes and rest. During the following fMRI scan, participants performed the reward task concurrent with pupillometric data collection (not reported here). After the fMRI scan, a neuromelanin-sensitive structural scan was acquired to assess LC integrity (not reported here).

Following the structural scans, participants performed the ‘immediate’ memory test for approximately 20 minutes outside the scanner (Figure 2). Subsequently, after a break, they performed a ‘delayed’ memory test, also lasting for about 20 minutes and conducted outside the scanner, at approximately 120 minutes post-reward task. During their second visit, participants were explicitly instructed not to engage in deliberate memorisation of the presented scenes to minimise the strategy effects in memory performance. Each memory test included a total of 176 items: 88 ‘old’ items, randomly selected from those presented during the incidental encoding reward task, and 88 ‘new’ items. The discrepancy in the number of trials between the encoding and recognition tasks was due to a limitation in the availability of new scenes to match the old items. This resulted in the random exclusion of four stimuli per subject presented during encoding from subsequent memory analyses. Among the old items, 66 were from the frequently presented category and 22 from the infrequently presented category. Similarly, the new items were also divided into 66 frequent and 22 infrequent scenes based on their scene category in order to prevent a bias in stimulus category frequency when comparing old and new scenes. Participants indicated whether a stimulus was old or new, as well as how confident they were in their assessment (‘sure’ or ‘not sure’) (Figure 2d). Pupillometric recordings (not reported here) were also acquired during the memory tests.

**Figure 2.**
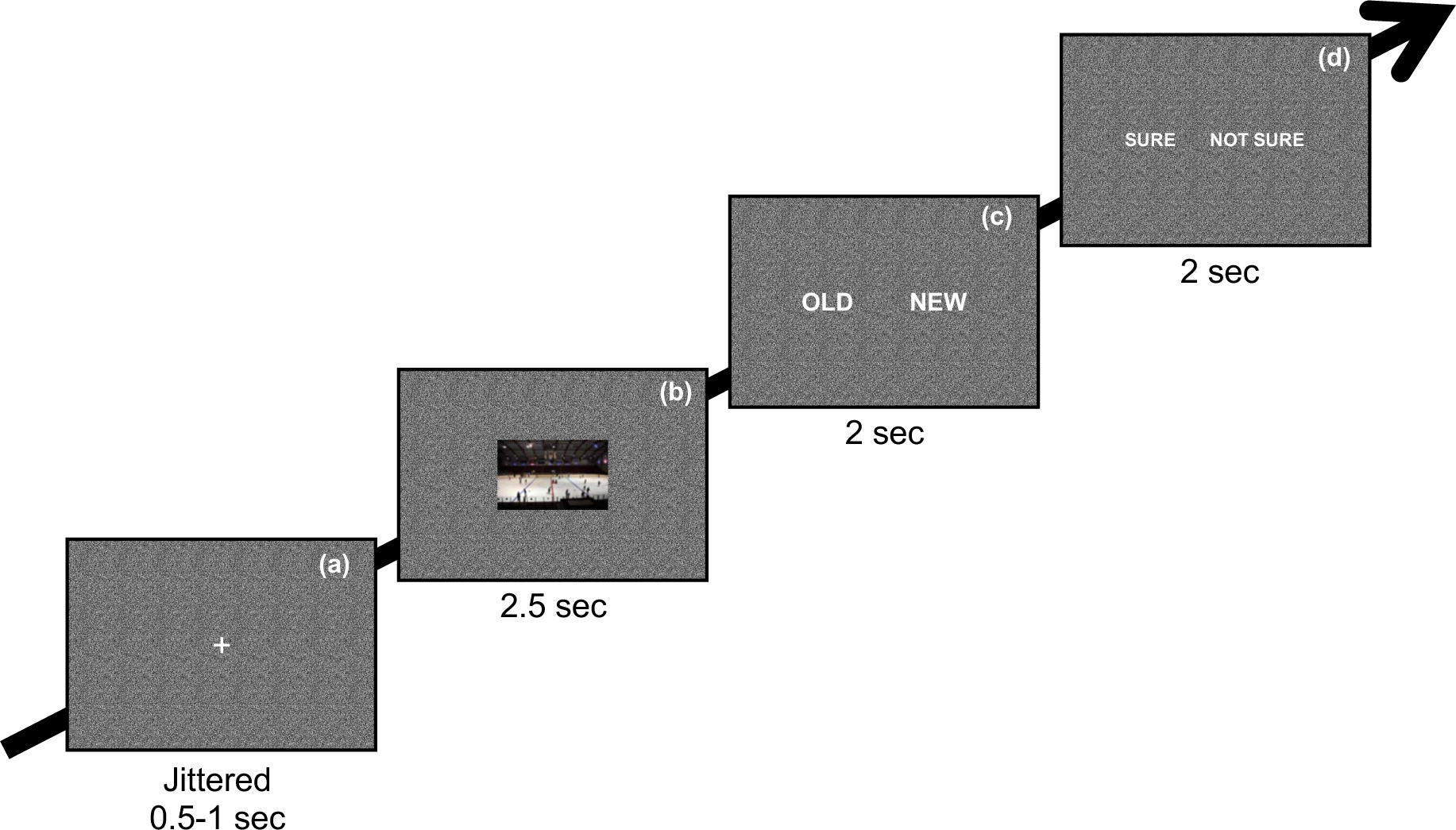
Incidental memory tests. The layout of the stimulus on the screen and the sequence within a trial: (a) baseline; (b) a scene which were either already seen during the reward task in the scan session or new; (c) an old-new recognition response in which participants were to respond whether they have seen the stimulus or not; (d) a binary confidence rating screen in which participants were to respond whether they are sure of their decision they made in the recognition response.

#### 2.2.3 Imaging protocols

All images were acquired with a Siemens 3T Biograph mMR scanner (Siemens Healthineers, Erlangen, Germany) using a 24-channel head coil.

##### 2.2.3.1 Structural MRI acquisition

Per session, a high-resolution T1-weighted anatomical image (MPRAGE) was acquired to support functional image co-registration (1mm isotropic voxel size, 192 slices, TR=2,500ms, TE=4.37ms, TI=1100ms, FOV=256×256×192mm, flip angle[FA]=7°), a coronally oriented T2 image to assess hippocampal subfield volumes (0.4×0.4×2mm voxel size, 29 slices, TR=8020ms, TE=52ms, FOV=175×175×58mm; not reported here), and an axially oriented high-resolution neuromelanin-sensitive T1-weighted multi-echo FLASH sequence to characterise LC integrity (0.6×0.6×3mm voxel size, 48 slices, TR=22ms, TE=5.57ms, TA=4:37, FOV=230×230×144mm, FA=23°; not reported here).

##### 2.2.3.2 Functional MRI acquisition

During the reward task, a T2*-weighted 3D EPI was acquired perpendicularly to the back of the brainstem (2mm isotropic voxel size, 51 slices, TR=3600ms, TE=32ms, FOV=240×240×102mm, FA=80°).

#### 2.2.4 Data preprocessing and analysis

##### 2.2.4.1 sMRI data

Individual T1-weighted whole-brain structural images underwent bias correction using the advanced normalization tool’s *N4BiasFieldCorrection* function (ANTs, Version 2.3.1). This correction was necessary to address field-related inhomogeneity in the images, which can hinder the normalisation of the images into the group space. The Montreal Neurological Institute (MNI) template space was used as the group space (38). A study-specific template space was created from these bias-field-corrected structural whole-brain images using *antsMultivariateTemplateConstruction2* function of ANTs (only one of the two T1w images collected per participant was selected) to allow for a more precise normalisation into group space. Parameters for bias correction and template generation are shown in the Supplementary Method 1.

##### 2.2.4.2 fMRI data

For each participant, functional scans from the two sessions underwent separate slice-time correction, and un-warping was performed using the respective field maps with Statistical Parametric Mapping (SPM12, http://www.fil.ion.ucl.ac.uk /spm12.html) within the MATLAB environment (Version 2015a, MathWorks, Sherborn, MA, USA, 2015) using default parameters. Subsequently, the scans from both sessions were concatenated and realigned using the default parameters of SPM12’s *Realign* functions to compare the frequent- and infrequent-reward conditions across sessions. Alignment quality was visually assessed. Functional scans were then smoothed with a 3×3×3mm kernel using SPM12’s *Smoothe* function, followed by single-subject voxelwise general linear model (GLM) analyses to estimate task-related contrasts in SPM12. Due to technical issues preventing physiological noise parameters from being recorded for 24 datasets, CompCor was applied uniformly during single-subject GLM analyses for consistency. This method has been shown to provide comparable results to regressor-based noise correction (39). The resulting contrast maps were transformed into the structural MNI template space for group analyses using a pipeline combining ANTs and FSL (FMRIB Software Library, Version 6.0.4). More details about the pipeline can be found in Supplementary Method 1.

##### 2.2.4.3 Quality assessment of the functional image transformation

To ensure that sufficient spatial precision was achieved in the transformation of individual data to the group space, quality assessments were conducted (YY), as described in Yi et al. (2023). Briefly, anatomical landmarks on the brainstem were delineated on each MNI-transformed mean functional image and compared to the corresponding landmarks on the structural MNI template. The spatial deviations between individual and pre-set landmarks were then calculated per participant and per landmark and were summarised across participants. As can be seen in Figure 3, deviations generally stayed below 2mm indicating sufficient precision in spatial transformations in the midbrain and brainstem.

**Figure 3.**
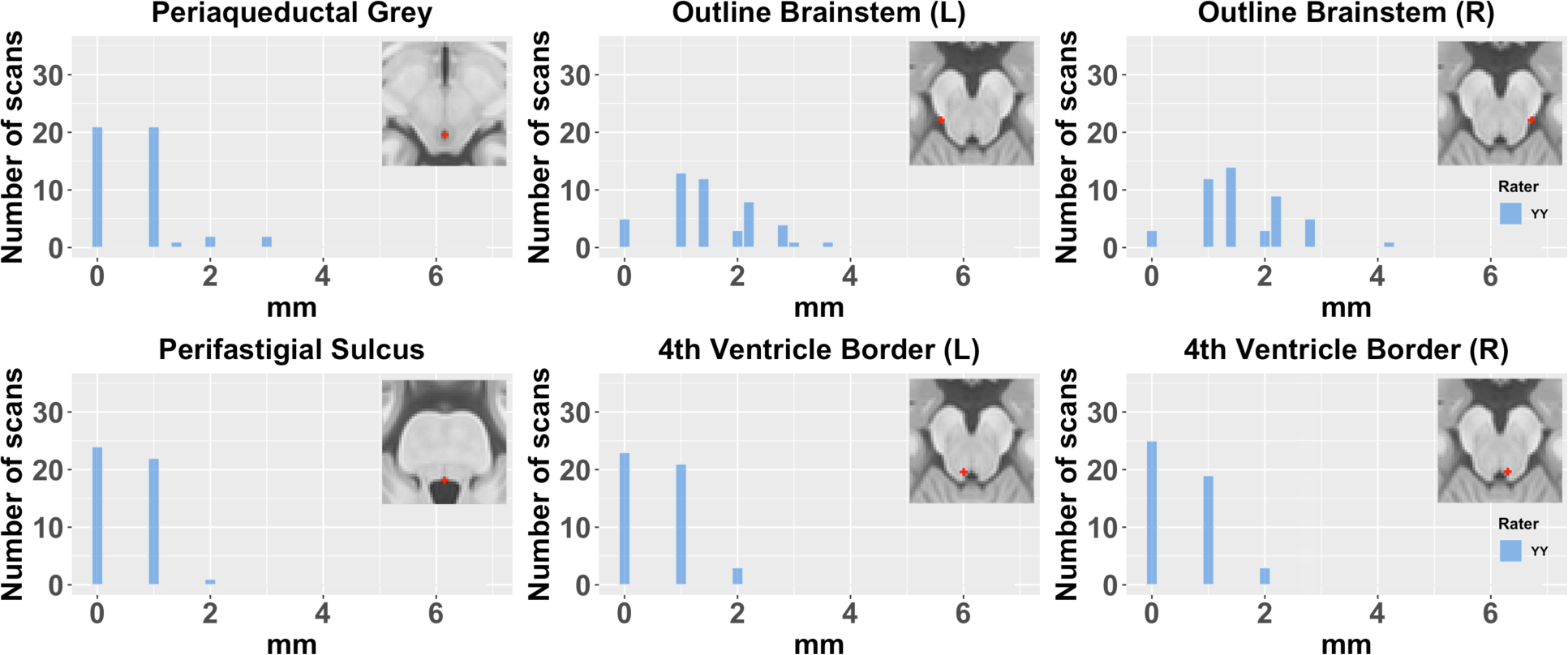
Histograms of in-plane distances between landmarks defined on the MNI template and single-subject landmarks delineated on MNI-transformed mean functional images. Each inset in the corresponding histogram plot indicates its anatomical position on the MNI template. The detailed procedure for selecting and placing the landmarks, as well as quantifying the distances, is described in Yi et al.’s (2023) work and Supplementary Method 2. Note that the distances in the Outline Brainstem landmarks vary, as they were placed anywhere along the outline of the brainstem border. The mean±standard deviation distances for landmarks are as follows: Periaqueductal Grey (0.69±0.76), Perifastigial Sulcus (0.51±0.55), Left Outline Brainstem (1.53±0.85), Right Outline Brainstem (1.62±0.82), Left 4th Ventricle Border (0.57±0.62), and Right 4th Ventricle Border (0.53±0.62).

##### 2.2.4.4 Masks and significance thresholds used in fMRI analyses

For whole-brain analyses, an inclusive grey matter mask segmented from the structural MNI template using the *Segment* function of SPM12 applied at *p*_uncorr_<.001 threshold was used. In these analyses, cluster-level significance was determined by applying the False Discovery Rate (FDR) method for multiple comparisons correction within the same *p*_uncorr_<.001 significance threshold, as per the approach outlined by Genovese, Lazar, & Nichols (40). An anatomical midbrain and brainstem mask was applied as an inclusive mask at *p*_uncorr_<.001 to investigate the small structures in the midbrain and brainstem (41). SN activation was examined with small-volume correction (SVC) with the SN mask extracted from Pauli et al.’s reinforcement learning atlas (42).

##### 2.2.4.5 Behavioural data

Behavioural data were analysed using SPSS (version 29, SPSS Inc., Armonk, NY, USA, 2021). To quantify memory performance under each condition (immediate/delayed tests, reward/neutral outcome, and infrequent/frequent presentation), the D-prime (*D’*) measure was computed. This metric was derived by first calculating the hit rate (*H*) and false-alarm rate (*F*) for each condition, with small corrections applied to prevent extreme values as outlined in Hautus (1995):

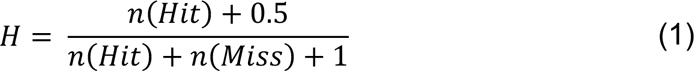

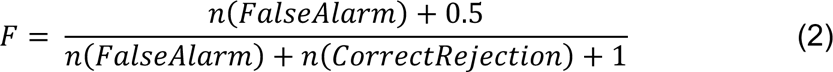

The *D’* values were then derived as the difference between the inverse cumulative distribution functions (Φ^−1^) of the corrected hit and false-alarm rates:

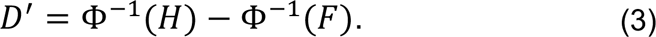

## 3 Results

As outlined previously, our task was designed to manipulate two distinct aspects of stimulus salience in two separate sessions: (a) the association of a stimulus with a reward versus a neutral outcome, referred to as “reward salience,” and (b) the association of a stimulus with a less frequent outcome, referred to as “contextual unexpectedness salience”. In the following analyses, we aimed to identify brain regions specifically associated with these two aspects of salience (i.e., reward and contextual unexpectedness). All fMRI GLM results were analysed using SPM12 in the MATLAB environment (version 2021a, Mathworks, Sherborn, MA, USA, 2021). A comprehensive list of all activations, their statistical significance, and their coordinates in Talairach space can be found in Supplementary Table 4 and 5.

### 3.1 Behavioural results

Participants exhibited a high accuracy of categorising the stimulus sets during the reward task in both infrequent and frequent reward sessions, with an average accuracy of 94% (*SD*=8%). A one-way ANOVA analysis showed no significant difference in categorisation accuracy between the two sessions, *F*(1,92)=0.642, *p*=.425. The results of the two-way ANOVA indicated no significant main effects of contextual unexpectedness (infrequent/frequent; *F*[1,184]=1.912, *p*=.168) or reward (reward/neutral; *F*[1,184]=1.576, *p*=.211) on the categorisation accuracy. In addition, there was no significant interaction between frequency and reward variables, *F*(1,184)=2.643, *p*=.106. Also, there was no significant main effects of delay length (immediate, *F*[1,92]=0.024, *p*=.877; delayed, *F*[1,88]=0.069, *p*=.793), reward (reward, *F*[1,88]=0.285, *p*=.595; neutral, *F*[1,88]=0.086, *p*=.690), and frequency (infrequent, *F*[1,88]=0.160, *p*=.690; frequent, *F*[1,88]=0.022, *p*=.883) on the memory test performances between the first and the second visit.

#### 3.1.1 Memory test performance

As outlined above, stimulus categories were counterbalanced across salience conditions. Memory performance across the four stimulus categories, did not differ (urban and nature from the outdoor category and private and public from the indoor category; One-way ANOVA, immediate memory test: *F*(3,183)=1.854, *p*=.139; delayed memory test: *F*(3,173)=2.074, *p*=.105).

To assess memory effects related to salience types, a three-factor repeated measures ANOVA was calculated (contextual unexpectedness [infrequent/frequent] × reward [reward/neutral] × delay length [immediate/delayed]) on D’. As expected, memory performance was higher for the immediate memory test as compared to the delayed memory test, *F*(1,42)=110.183, *p*<.001, as well as for infrequently presented scenes compared to frequently presented scenes, *F*(1,42)=21.954, *p*<.001. The better memory for infrequently presented scenes is in line with previous studies showing an association between unexpected or contextually salient events and improved recollection performance (von Restorff or isolation effect; 26–28,44–46). Moreover, a significant interaction effect between contextual unexpectedness and delay length factors, *F*(1,42)=21.181, *p*<.001, η_p_^2^=.335, indicates that the contextual unexpectedness effect was more pronounced on the immediate memory test. This suggests that the advantage of stimulus salience for memory is most prominent in the short-term and may not persist over longer periods if the stimulus’ episodic salience is less pronounced (21,26,31,47).

In addition, a three-factor repeated measures ANOVA (contextual unexpectedness [infrequent/frequent] × reward [reward/neutral] × delay length [immediate/delayed]) performed on the memory tests’ reaction times (RTs) showed faster RT to infrequently presented scenes than to frequently presented scenes, *F*(1,42)=6.962, *p*=.012, suggesting also stronger memory traces for infrequently presented scenes (44,48–51). Similarly, slower RTs during the immediate memory test than delayed memory test were observed, *F*(1,42)=25.204, *p*<.001, which might imply that scenes that had formed stronger memory traces form a more prominent portion of the old responses in the delayed test (51,52). A trend of an interaction between contextual unexpectedness and delay length showed slightly faster RTs for infrequently presented scenes during the delayed memory test than the immediate memory test, while RTs for frequently presented scenes remain unchanged across the two memory tests, *F*(1,42)=3.082, *p*=.086, η_p_^2^=.068, no two- or three-way interaction effect among the factors was found. Likewise, this trend in RT performance likely indicates that infrequently presented scenes may have been encoded more robustly (52,53).

Unexpectedly, there was no memory effect for reward-associated as compared to neutral scenes, indicating a comparatively weaker memory-relevant effect of reward salience in our setup for combining unexpected and rewarded events, *F*(1,42)=2.229, *p*=.143 (Figure 4). The observed lack of a significant memory enhancement for rewarded compared to non-rewarded scenes could be attributed to several factors, not all of which are mutually exclusive. First, to avoid diverting attention from the unexpectedness of rare stimuli in the infrequent stimulus category, reward feedback was deterministically and not probabilistically related to reward scenes. However, previous research suggests that probabilistic rewards generate larger reward prediction errors (RPEs) (54–56), a potential enhancement to memory effects that our deterministic approach might not have fully captured. Moreover, it has been suggested that associations with rewards have a stronger effect on decision biases, namely, a bias towards approaching stimuli rather than enhancing memory discrimination (57).

**Figure 4.**
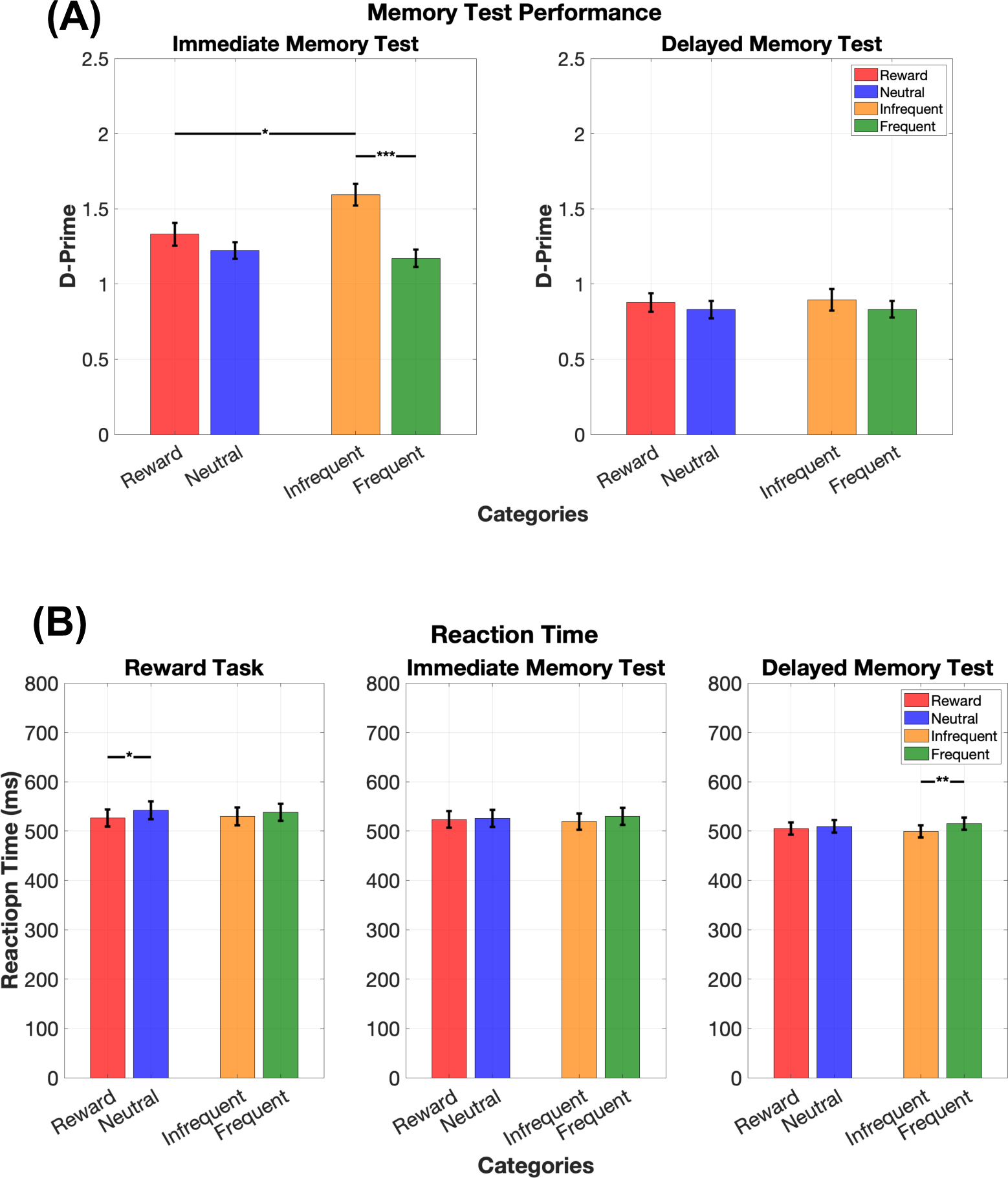
Memory test performance in immediate and delayed recognition tasks and reaction time (RT) performance during the reward task and immediate and delayed recognition tasks for the two salience manipulations. (A) displays the D’ results for the immediate (left) and delayed (right) memory tests, **encompassing all trials**. Each bar plot from left to right represents the D’ values for scenes associated with reward, neutral, infrequently presented (infrequent), and frequently presented (frequent) scenes. **(B)** represents the RT performance in response to prompts (scene category judgment [e.g., private vs. public] during the reward task and [old vs. new] during recognition memory tests), which were presented following a scene stimulus. In both top and bottom panels, horizontal bars with asterisks denote significant differences between stimulus categories. One asterisk (*) represents *p*<0.05, and three asterisks (***) represent p<0.001 significance threshold.

Specifically, Bowen et al. (57) observed that although reward-associated stimuli can increase hit rates, this did not translate into an increased *D’*. The authors explain that this phenomenon may arise from reward salience primarily influencing decision-making tendencies, leading to a more liberal response bias towards stimuli associated with rewards during recognition tests. Indeed, in our results, although participants showed better recognition of familiar reward-associated scenes (Supplementary Figure 3C and 3D), this was offset by a larger increase in FA for these scenes (Supplementary Figure 3A and 3B), resulting in no overall change in D’. This result is similar to what was found in Bowen et al. (57), who employed a similar encoding task paradigm (Experiment 1) as this study, and demonstrated that high-reward cues increase hit rates without necessarily enhancing memory discriminability (*D’*). This suggests that reward motivation affects decision biases rather than memory discrimination. This leads to a more liberal response bias in recognition tests (57), resulting in increased rates of both hits and false alarms (Supplementary Figure 3). Corroborating this, although no significant differences in RTs were observed between frequent and infrequent stimuli during the encoding, RTs were significantly quicker for scenes associated with rewards compared to neutral ones, *F*(1,46)=5.448, *p*=.024. This is in line with prior studies demonstrating faster RTs when approaching reward-associated stimuli (‘action vigor’; 56,57).

When restricting the analysis to high-confidence trials to assess items with stronger memory traces, results paralleled those observed in the full trial set. There was a main effect of contextual unexpectedness, *F*(1,42)=16.740, *p*<.001, and delay length, *F*(1,42)=82.260, *p*<.001, along with an interaction effect between these factors, *F*(1,42)=10.150, *p*=.003, η_p_^2^=.195, further confirming a robust effect of contextual unexpectedness and delay length on memory.

To explore the impact of salience types on false alarms (FAs), a three-factor repeated measures ANOVA was conducted. Main effects showed higher FA in delayed than immediate tests, consistent with the generally weaker memory performance on delayed tests, *F*(1,42)=16.309, *p*<.001. However, no significant differences were found for reward or contextual unexpectedness. Significant two-way interactions were observed between delay length and both reward and contextual unexpectedness on FAs (Supplementary Figure 3A and 3B). Specifically, both reward-associated and neutral scenes initially showed similar FAs during the immediate memory tests. However, reward-associated scenes exhibited a sharper increase in FAs compared to neutral scenes with longer delays (Supplementary Figure 3A), *F*(1,42)=4.137, *p*=.048, η_p_^2^=.090. In contrast, although there was a trend in the main effect of contextual unexpectedness showing that infrequently presented scenes had lower FAs compared to frequently presented ones, *F*(1,42)=3.839, *p*=.057, infrequently presented scenes showed an increase in FAs in delayed memory tests, while the FAs for frequently presented scenes remained largely unchanged (Supplementary Figure 3B), *F*(1,42)=6.995, *p*=.011, η_p_^2^=.143. These results indicate their differential effects of salience types on FA over time. However, no interaction between reward and unexpectedness or any three-way interaction was observed. These interactions suggest that the temporal delay between encoding and recognition modulates FAs in a salience-dependent manner. Yet, there were no significant interactions between reward and unexpectedness, nor any three-way interaction, highlighting that salience types alone may not differentially affect FAs.

Regarding hit-rate analyses, as expected, a three-factor repeated measures ANOVA revealed higher hit rates for immediate than delayed memory test, *F*(1,42)=108.992, *p*<.001. A significant main effect of reward was also observed, *F*(1,42)=19.829, *p*<.001, indicating that hit rates were higher for reward-associated scenes than for neutral scenes. Additionally, a smaller, yet significant main effect of contextual unexpectedness was found, *F*(1,42)=10.360, *p*=.002, showing higher hit rates for infrequently presented scenes. As for interaction effects, the interaction between the delay length and reward exhibited a trend (Supplementary Figure 3C), *F*(1,42)=3.711, *p*=.061, η_p_^2^=.081, suggesting an initially nonsignificant effect of reward on hit rate in the immediate memory test becoming more pronounced in the delayed memory test. The interaction between delay length and contextual unexpectedness was also significant (Supplementary Figure 3D), *F*(1,42)=6.088, *p*=.018, η_p_^2^=.127, indicating that the initial advantage in the hit rate due to contextual unexpectedness during the immediate memory test did not persist into the delayed memory test.

#### 3.1.2 Confidence ratings during immediate and delayed memory tests

Binary confidence ratings (0 – ‘not sure’, 1 – ‘sure’) were averaged within each of the four conditions (contextual unexpectedness [infrequent/frequent] × reward [reward/neutral]) and separately for correct (hit and correct rejection) and incorrect (FA and miss) trials on the memory tests. Two three-factor repeated measures ANOVA found that, in *correct trials*, confidence ratings were higher to infrequently presented items than frequently presented items, *F*(1,42)=31.261, *p*<.001, and higher in immediate memory test than delayed memory test, *F*(1,42)=23.410, *p*<.001. However, no significant reward effect was found, and there was no interaction effect across all variables. In *incorrect trials*, only immediate memory tests showed higher confidence ratings than delayed memory tests, *F*(1,42)=6.686, *p*=.013. This effect in delay length (immediate/delayed) suggests a possible recency effect, where participants may feel more confident about their answers in an immediate memory test because the information is still relatively fresh in their minds, even if they are incorrect (60).

In summary, our findings align with the von Restorff effect (26–28,44–46), showing that varying contextual unexpectedness as a form of salience manipulation consistently influences memory performance. Specifically, scenes categorised as ‘infrequently presented’ were better remembered than those in the ‘frequently presented’ category This effect was particularly pronounced in immediate memory tests, where the impact of contextual manipulation was more present, as the encoding context is comparatively more recent and most similar to the retrieval context 27/05/2024 18:34:00. Furthermore, faster RTs associated with ‘infrequently presented’ scenes during memory tests may indicate stronger memory traces for these infrequent stimuli, an effect that was especially marked in delayed memory tests.

Contrary to expectations, reward-associated scenes did not show enhanced memory effects compared to neutral scenes. This could be due to a) the use of deterministic feedback resulting in a potentially weaker reward manipulation, and b) reward associations having a more significant impact on decision biases than memory discrimination (57). It is important to note that this does not imply reward associations had no effect on a differential processing of rewarded versus non-rewarded stimuli. Indeed, we observed shorter RTs to reward-associated scenes during encoding, in line with previous studies that observed faster RTs towards reward-associated stimuli (58,59). Moreover, although the ratio of hits to FAs remained unchanged between rewarded and neutral scenes, scenes from the reward-associated category were more frequently classified as ‘old’ during memory tests compared to neutral scenes. This suggests a greater inclination to perceive reward-associated stimuli as familiar, again indicating reward-influenced decision biases.

While our results suggest a stronger effect of contextual unexpectedness on memory processes, reward associations therefore still yielded expected effects for rewarded stimuli, albeit more in the domain of affecting decision biases and RTs in favour of reward-associated stimuli. These differential effects of saliency manipulations, reward and contextual unexpectedness, are interesting in their own regard. However, they also pose challenges in directly comparing their impact within our experimental paradigm. In the following we therefore focus in particular on a qualitative rather than a quantitative comparison of brain processes underlying the two salience manipulations.

### 3.2 fMRI results

In examining the fMRI data, we aim to assess whether two types of salience, as defined by reward and contextual unexpectedness, elicits differential activation, particularly within the midbrain and brainstem regions. Drawing from previous research involving both human and animal subjects, we hypothesised that reward-associated salience and memory would engage midbrain dopaminergic nuclei SN and VTA (63), subcortical areas such as the nucleus accumbens (64), amygdala (65,66), and other components of basal ganglia such as caudate and putamen (67), and cortical areas such as insular cortex (68,69) and orbitofrontal cortex (67,70). On the other hand, infrequent or contextually unexpected events would preferentially engage brainstem nuclei, such as the locus coeruleus (71,72). However, co-activation of the SN and VTA (10,31,73) may also occur. We further predicted that subcortical and cortical areas from the salience network, including amygdala (65,66), the inferior, medial, and superior frontal gyri (65,74–76), the temporoparietal cortex (65,77), and the anterior cingulate cortex (ACC; 67,70) would be additionally engaged during the processing and memory encoding of unexpected events.

For a detailed information on the model specifications and GLM contrasts utilised in our fMRI analyses, please refer to Supplementary Tables 1, 2, and 3, which outline predictor properties, contrast coding, and control predictors employed in the first-level models as described in sections 3.2.1 to 3.2.3. Also, a comprehensive list of fMRI activations can be found in Supplementary Table 4 and 5.

#### 3.2.1 Infrequently presented trials vs. frequently presented trials

##### 3.2.1.1 Scene presentation timepoint

As can be seen in Figure 5A (in green to yellow shade), bilateral insular cortex, bilateral parahippocampal gyrus (PHG), bilateral ventromedial caudate (the head of caudate), bilateral inferior parietal lobe, right ACC were more engaged during scenes from the infrequently presented scene categories. As also outlined above, the insular cortex, inferior parietal lobe, and ACC are known components of the salience detection and attentional modulation network (78–81). In addition, the observed bilateral ventromedial caudate activation may suggest inputs from the SN, as supported by histology and connectivity studies (82).

**Figure 5.**
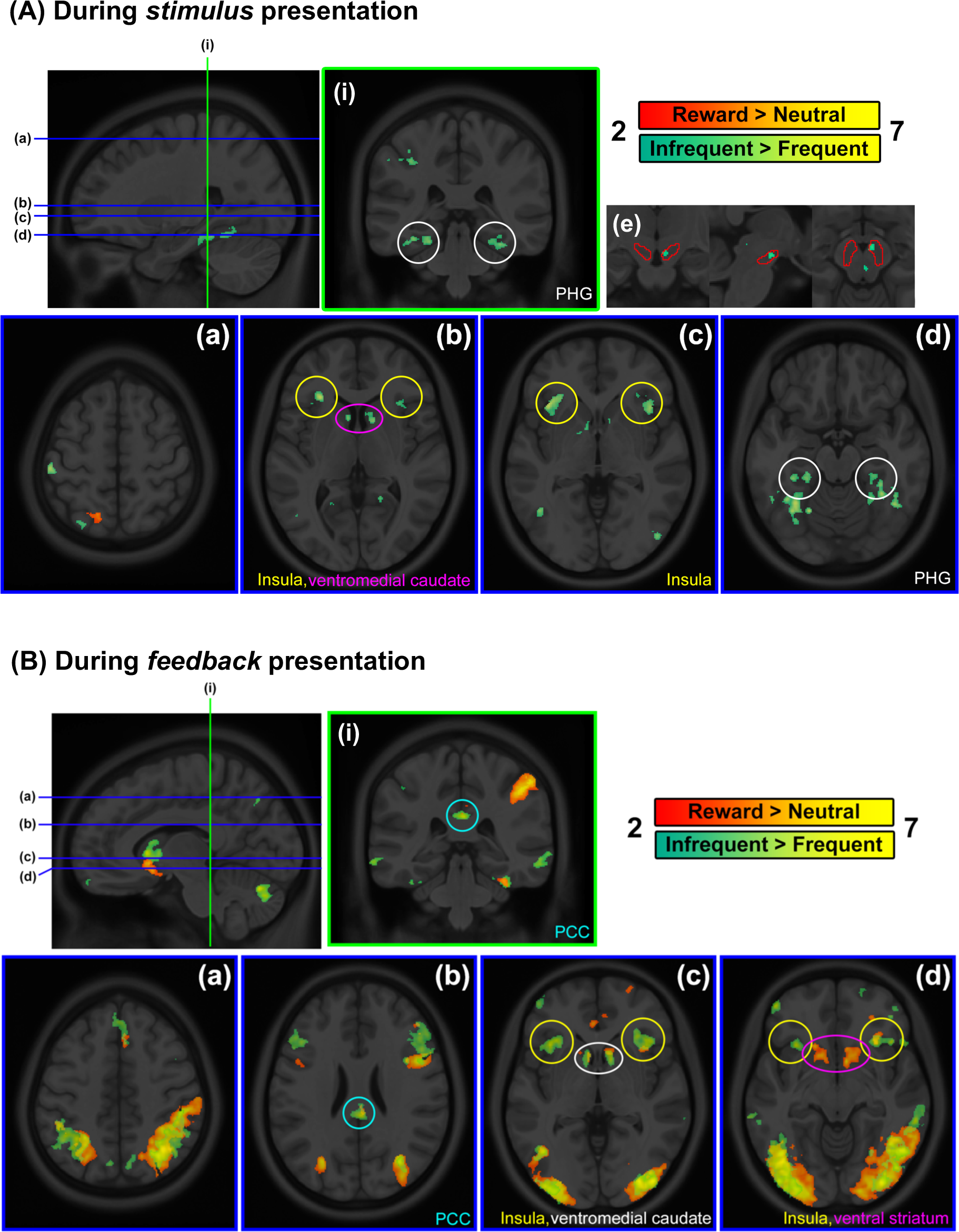
fMRI results from the infrequently versus frequently presented categories and the reward versus neutral categories. All activations were found with significance threshold of *p*_uncorr_<.001 and was FDR-controlled except for small-volume correction (SVC) analysis, which was examined with significance threshold of *p*_uncorr_<.001 but not FDR-controlled. **(A) Activations during scene presentation**: For activations during **reward-associated scene presentation**, axial slice (a) shows activation in the left superior parietal lobule compared to neutral trials. For activations during **infrequently presented scene presentation**, axial slice (b) and (c) demonstrate bilateral activation in the anterior caudate and insula, respectively, while axial slice (d) and coronal slice (i) display bilateral activation in the parahippocampal gyrus (PHG) compared to frequently presented scenes. Insets (e) show the right *dorsal* SN activation (SN mask used for SVC is delineated with red lines. X=6, y=-14, z=-14; Z_E_=4.15; *p*_FWEc_<0.05, *k*_E_=29). **(B) Activations during feedback presentation**: Axial slice (a) shows bilateral medial superior frontal cortex; (c) shows bilateral ventromedial caudate and insula activation; and axial slice (b) and coronal slice (i) show bilateral posterior cingulate cortex (PCC) activation in **infrequently presented feedbacks** compared to frequently presented feedbacks. In **reward-associated feedbacks** compared to neutral feedbacks, activation profiles mostly overlap, except, as seen in the axial slice (d) and sagittal slice, a bilateral ventral striatum activation is observed in comparison to bilateral ventromedial caudate activation in infrequently presented versus frequently presented feedbacks contrast.

Using the inclusive midbrain and brainstem mask to focus specifically on neuromodulatory nuclei in the brainstem, we furthermore observed higher right SN activation for infrequently presented scenes in the midbrain (small-volume corrected [SVC], x=6, y=-14, z=-14; *Z_E_*=4.15; *p*_FWEc_<0.05, *k*_E_=29, Figure 5A, the top right figure set). This is well in line with studies showing higher SN activations to novel or unexpected events (31,83,84). On the other hand, no significant activation was observed in the brainstem.

##### 3.2.1.2 Feedback presentation timepoint

During feedback presentation, several regions showed significant activation, including the insular cortex, inferior parietal lobule, ventromedial caudate, and posterior cingulate cortex (PCC) among others (see Figure 5B). These activated regions are reported to be associated with several cognitive functions such as attentional control (78,85), and reward processing (86,87). No significant activation was observed in the midbrain and brainstem.

Taken together, the processing of unexpected stimuli appears to be partly supported by the dopaminergic system. This is evidenced by the activation of SN, typically linked to dopamine, together with likely target regions such as the ventromedial caudate. The higher activations in cortical areas such as ACC, PCC, and insular cortex were expected as these structures are part of the salience network (66).

#### 3.2.2 Reward trials vs. neutral trials

##### 3.2.2.1 Scene presentation timepoint

On the whole-brain level, the left superior parietal lobe showed stronger activation for reward-associated scenes (Figure 5A, in red to yellow shade). No significant cluster was found in the midbrain and brainstem.

##### 3.2.2.2 Feedback presentation timepoint

On the whole-brain level, bilateral middle occipital lobes, bilateral anterior insular cortex, bilateral ACC, bilateral nucleus accumbens, bilateral ventromedial caudate, right middle cingulate cortex (MCC), and left inferior temporal lobe (ITL) showed stronger activation for reward feedback as compared to neutral feedback (Figure 5B in red to yellow shade). This activation pattern in anterior insular cortex, ACC, ventromedial caudate, and nucleus accumbens is corroborated by previous studies that investigated attentional control and reward assessment (78,79,88–90).

It should be noted that, as mentioned in the memory test performance of reward-associated scenes (item 3.1.1), the absence of activation in midbrain regions associated with reward salience, such as the SN or VTA during feedback might be attributed to the absence of RPEs. As our task aimed at orthogonally modulating reward salience and contextual unexpectedness, reward feedbacks were deterministically followed by reward-associated scenes, resulting in reward processing without prediction errors. These weaker reward-related responses may have resulted in weaker responses in these areas typically implicated in reward processing (55,91,92).

A comprehensive list of activation clusters and statistical results of each cluster from this contrast can be found in the Supplementary Table 4.

#### 3.2.3 Subsequent memory effects

In the subsequent-memory analysis, only hits, i.e., items correctly identified as old, were included from both immediate and delayed memory tests, which were pooled together. To isolate the effect of the two saliency types on memory encoding, scene stimulus presentation timepoints were analysed. This approach minimises potential confounding variability introduced by reward feedback, which, while informative, is already anticipated by subjects due to pre-task conditioning. Details of the GLM model predictors and contrast coding configuration regarding the analyses included in this item are delineated in Supplementary Table 2 and 3. We will first assess which areas are more activated for remembered salient scenes compared to remembered non-salient scenes, to investigate which brain areas distinguish stimulus salience during memory encoding (3.2.3.1 and 3.2.3.2). This will be followed by a 2×2×2 comparison of the two salience effects on memory, where we will examine the joint effects of reward and contextual salience on memory enhancement (3.2.3.3, cf. Supplementary Table 5). Finally, we examined memory-specific processes separately for each salient stimulus category by contrasting remembered and forgotten scenes within each type, aiming to identify brain areas that support the memory formation for salient stimuli, the results of which can be found in Supplementary Results 3 and Supplementary Figure 5.

##### 3.2.3.1 Subsequently remembered infrequently presented vs frequently presented scenes

During the *scene* presentation, subsequently remembered ***infrequently presented*** scenes as compared to remembered *frequently presented* scenes showed greater activation in the left calcarine sulcus, left precuneus, bilateral postcentral gyrus, right inferior frontal cortex, left inferior parietal lobe, left fusiform gyrus, and left superior medial frontal cortex (Figure 6). This supports the idea that these areas, which are involved in visual and semantic processing (calcarine sulcus and inferior parietal lobe: (93,94)), retrieval and integration of memory (precuneus: (95)), and attentional control and monitoring of memory processes (the inferior frontal gyrus and superior medial frontal gyrus: (96)), are more engaged during the encoding and retrieval of the salient, infrequently presented scenes. Importantly, a significant activation in the right dorsal SN was found for these better remembered infrequently presented scenes, suggesting that the encoding of scenes associated with unexpectedness-related salience is likely associated with dopaminergic activity in the SN (SVC; x=6, y=-15, z=-14; *Z*_E_=4.15; *p*_FWEc_<0.05, *k*_E_=31).

**Figure 6.**
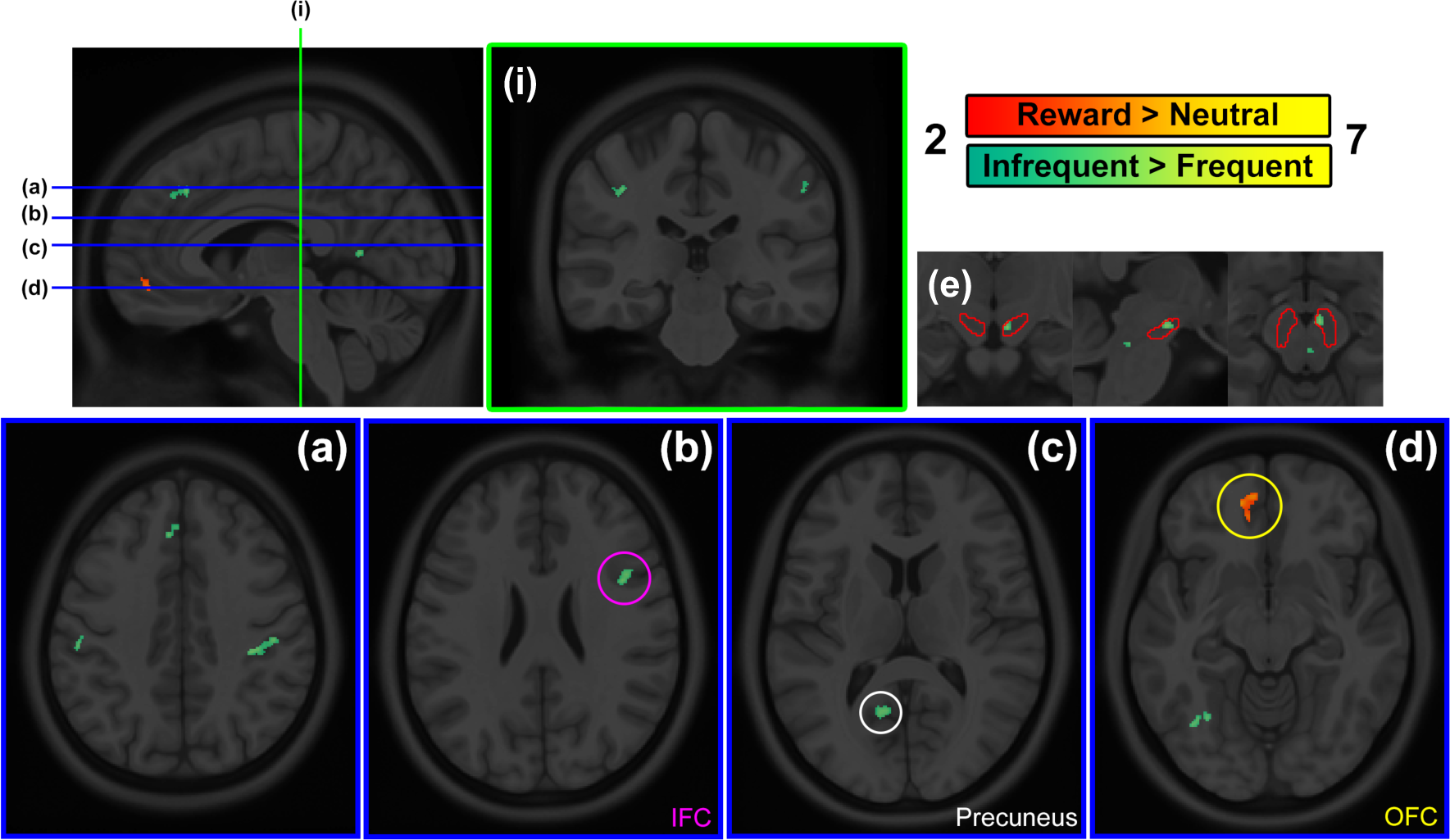
fMRI results from the infrequent versus frequent scenes and the reward versus neutral scenes in the subsequently remembered scenes. All activations were found with significance threshold of *p*_uncorr_<.001 and was FDR-controlled except SVC analysis, which was examined with significance threshold of *p*_uncorr_<.001 but not FDR-controlled. In subsequently remembered *infrequently presented* scenes compared to *frequently* presented scenes, coronal slice (i) and axial slice (a) shows activations in bilateral postcentral gyrus and left superior frontal cortex (SFC); axial slice (b) shows right IFC; (c) shows left precuneus; and (d) shows left calcarine sulcus. On the other hand, during the presentation of subsequently remembered *reward-associated* compared to subsequently remembered *neutral* scenes, left orbitofrontal cortex (OFC) showed significant activation, as seen in sagittal slice and axial slice (d). As shown in insets (e), an SVC analysis on this contrast found right *dorsal* SN activation for subsequently remembered *infrequently presented* scenes compared to *frequently* presented scenes (SN mask used for SVC is delineated with red lines. X=6, y=-15, z=-14; *Z*_E_=4.15; *p*_FWEc_<0.05, *k*_E_=31).

The activation of frontal and parietal regions might indicate an additional involvement in enhanced visual processing and attention, in line with prior research implicating these regions in memory tasks and visual perception (97–99).

##### 3.2.3.2 Subsequently remembered reward-associated vs neutral scenes

When comparing reward-associated scenes that are subsequently remembered versus subsequently remembered neutral scenes, only the left orbitofrontal cortex was more activated (Figure 6). This suggests that the reward-related information was better encoded and consolidated, which led to better retrieval of the memory during the recognition phase of the task. This could be related to the role of the region in evaluating the reward value of stimuli and guiding behaviour accordingly (70,100).

##### 3.2.3.3. Interaction among contextual unexpectedness, reward, and memory

In our examination of the mechanisms supporting the effect of contextual unexpectedness and reward on memory, we sought to understand how the different types of salience interact with each other to influence memory. To this end, we conducted a full factorial ANOVA focused on these three factors, contextual unexpectedness (infrequent > frequent), reward (reward > neutral), and memory outcome (remembered > forgotten) (Supplementary Table 2). Intriguingly, our analysis did not reveal any significant cortical activations for all inspected two- and three-way interaction pairs. However, an interesting dissociation in SN engagement was observed upon applying the inclusive midbrain and brainstem mask to inspect specifically on neuromodulatory nuclei in the brainstem. The left dorsal SN showed higher activation for infrequent and rewarded scenes, independent of memory outcome (SVC; [cluster 1: x=-8, y=-14, z=-13; *Z*_E_=4.51; *p*_FWEc_<0.05, *k*_E_=52], [cluster 2: x=-12, y=-19, z=-10; *Z*_E_=3.83; *p*_FWEc_<0.05, *k*_E_=35]), while the bilateral ventral SN was more activated for subsequently remembered infrequently presented scenes, independent of reward (SVC; [right: x=-7, y=-18, z=-19; *Z*_E_=3.75; *p*_FWEc_=0.06, *k*_E_=13], [left: x=8, y=-17, z=-16; *Z*_E_=3.93; *p*_FWEc_<0.05, *k*E=23]; Figure 7). No significant supracluster activation, either cortical or subcortical, was found in the three-way interaction among frequency, reward, and memory outcome.

**Figure 7.**
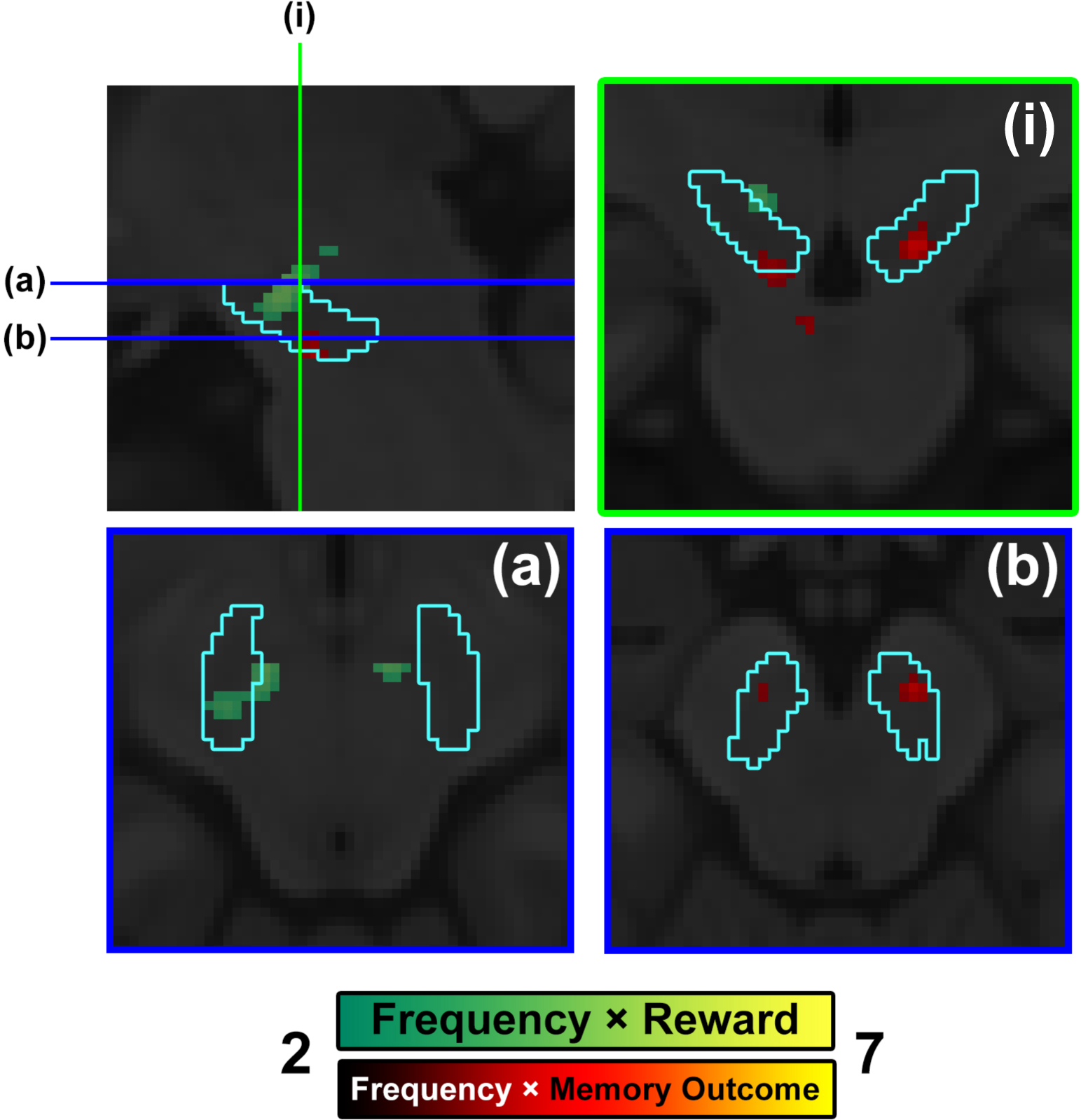
fMRI results from three-factor factorial ANOVA analysis testing positive interaction among contextual unexpectedness, reward, and memory. All activations were found with significance threshold of *p*_uncorr_<.001 within the inclusive brainstem mask and was not FDR-controlled. In the activation observed in **the positive interaction between Frequency (contextual unexpectedness) and Reward factors**, two clusters of activations in the left dorsal SN were found in an SVC analysis (sagittal, coronal, and axial slice [a]; [cluster 1: x=-8, y=-14, z=-13; *Z*_E_=4.51; *p*_FWEc_<0.05, *k*_E_=52], [cluster 2: x=-12, y=-19, z=-10; *Z*_E_=3.83; *p*_FWEc_<0.05, *k*_E_=35]). In **the positive interaction between Frequency and Memory outcome factors**, bilateral activations in ventral SN were found in an SVC analysis (sagittal, coronal, and axial slice [b]; [right: x=-7, y=-18, z=-19; *Z*_E_=3.75; *p*_FWEc_=0.06, *k*_E_=13], [left: x=8, y=-17, z=-16; *Z*_E_=3.93; *p*_FWEc_<0.05, *k*_E_=23]). SN mask used for SVC is delineated with cyan lines.

This subcortical emphasis in the SN highlights its important role in modulating the interactions between the salience of stimuli and their successful memory encoding. The significant activation observed within the right dorsal and ventral segments of the SN further implies the functional differentiation within the SN in encoding salience, aligning with documented functional heterogeneity that suggests a differentiated role of these SN subregions in modulating cognitive processes under varying reward conditions (101). These findings may indicate a specific dopaminergic mechanism within the SN that preferentially responds to the confluence of unexpectedness and reward, and their combined effect on successful encoding (102,103).

## 4 Discussion

In the present study, we aimed to investigate the impact of two types of salience, reward and contextual unexpectedness, in a 2-by-2 design on stimulus processing and incidental memory. As neuromodulatory nuclei of the midbrain and brainstem are important modulators of salience-related processing, we utilised high-resolution, high-precision fMRI recordings and analyses to investigate in particular the role of small subcortical nuclei in processing these two distinct types of salience.

Our behavioural findings revealed distinct effects of the two salience types on memory encoding and decision biases. Specifically, in line with the ‘von Restorff effect’ or isolation effect, which postulates better memory for contextually salient or unexpected events (33,34,44–46,104), memory performance was significantly enhanced for frequently presented scenes. This effect was particularly evident during immediate tests compared to delayed tests, suggesting that the advantage of stimulus salience may not persist over longer periods (21,26,31,47). Memory effects related to contextual unexpectedness were further confirmed by higher confidence ratings for infrequently presented items than for frequently presented items, in particular on immediate memory tests.

In contrast to the better subsequent memory for contextually unexpected scenes, scenes from reward-associated stimulus categories were not better remembered than those from neutral categories. However, reward associations still produced the typical reward-associated behavioural effects by affecting decision biases and RTs in favour of reward-associated stimuli. Specifically, we observed faster RTs for reward-associated scenes during the encoding task, along with heightened hit and FA responses to these scenes during memory tests, in line with previous reports of reward influencing ‘response vigor’ and decision biases (57–59).

Taken together, the behavioural results of our study suggest that contextual unexpectedness has a greater impact on memory processes as compared to reward association. Nevertheless, reward associations yielded expected effects, primarily manifesting in decision biases and response times favouring reward-associated stimuli. When comparing brain activations across the two salience types, these qualitative differences in associated processes thus need to be considered. We therefore focused on a qualitative rather than quantitative comparison of the brain mechanisms behind the two saliency modifications.

### 4.1. Distinct Brain Activation Patterns: Reward vs. Contextual Unexpectedness

In line with our expectations, distinct activation patterns for the two salience types were observed. For the reward versus neutral contrast, these were most notable at the feedback timepoints. In contrast, for the infrequent versus frequent scene stimuli, effects were pronounced both during the scene and feedback presentations. Given the deterministic association of stimulus categories with feedback, a stronger reward effect might have been expected already at the scene timepoints, consistent with studies showing reward cue effects (69). Nonetheless, feedback valence effects have been observed to persist even if feedbacks do not carry new information or are expected (105), suggesting that the mere exposure to desired or non-desired feedbacks remains emotionally and attentionally relevant, even without any new informational value.

Reward-associated feedbacks activated the nucleus accumbens, a central structure in the reward circuitry vital for processing reward, motivation, and reinforcement learning (106,107). Conversely, infrequently presented as compared to frequently presented scenes were most prominently accompanied by activations in the dorsal SN, insula, anterior caudate, and PHG. The anterior caudate, critical for integrating actions and outcomes (108–110), plays a critical role in enhancing visuo-motor associative learning, driven by phasic bursts of dopaminergic activity in response to unexpected events (110,111). This activity persists until the association is fully learned, maintaining elevated synaptic weights in caudate neurons as long as behavior is linked with the stimuli. Over time, as the learning consolidates, this activity gradually decreases (111). The larger activation for infrequently presented compared to frequently presented scenes is likely due to ongoing associative learning with infrequently appearing associations, whereas the frequent counterparts, having been sufficiently learned, show decreased activity levels. The PHG likely contributes to processing and encoding of contextually unexpected scene stimuli as it is known to be involved in novel information detection and encoding (112,113) and the processing of contextual associations (114) as well as the perception of visual scenes itself (115). Consistent with this finding, improved memory test performance, as indicated by *D’*, was observed in particular for contextually unexpected, or infrequent, stimuli.

Contrary to our expectations, we did not find the noradrenergic locus coeruleus to be involved in the processing of unexpected stimuli, despite our data acquisition protocols and analysis methods being specifically chosen to facilitate the identification of activations in small brainstem and midbrain nuclei. Given the smaller volume of the locus coeruleus compared to the SN, it is conceivable that larger sample sizes or longer acquisition durations than those included in our study would have been necessary. Nonetheless, our study was able to identify activations in subregions of the SN, which in volume are more similar to the locus coeruleus. Alternatively, it is possible that the paradigm employed was not ideally suited to evoke detectable changes in locus coeruleus activity given this sample size. As locus coeruleus imaging studies in humans are still sparse (35), it remains unclear whether results from animal studies suggesting an involvement of the LC in processing novelty or rewards (116) are easily translatable to the human domain. Indeed, a recent study observed larger LC activations during negative events and associated subsequently remembered stimuli, suggesting that negative stimulus valence might have stronger effects than unexpectedness (117). These limitations highlight the need for further, targeted research employing imaging with high signal-to-noise ratios in the brainstem and midbrain, and cognitive tasks with more robust manipulations of unexpectedness and valence.

Finally, our study suggests potential functional specialisations within the cingulate cortex for processing various salience types: MCC to reward, PCC to unexpectedness, and ACC to both (cf. Figures 5). This pattern might suggest distinct pathways and resource allocation strategies, contingent on salience type. The PCC and precuneus might have supported increased attention allocation to contextually unexpected events (118,119). Moreover, the co-activation of the insula and the ACC, both components of the salience network, appears to support processing of both reward and contextual unexpectedness (66,81,120,121).

### 4.2. Subcortical Modulation of Salience via SN and Its Effect on Memory Encoding

Intriguingly, we observed a distinction between the dorsal and ventral SN related to processing stimulus salience and the memory encoding of salient stimuli, respectively. Specifically, activations within the dorsal SN supported the processing of stimulus salience, as indicated by higher activity for infrequent compared to frequent scenes (cf. Figures 5, 7, and 8), as well as the interaction of infrequent larger than frequent and reward larger than neutral scenes (cf. Figure 8). Conversely, the bilateral ventral SN showed greater activation in processing salient (infrequent) scenes that were subsequently remembered (cf. Figure 8).

This distinction is in line with the evidence from studies documenting anatomical and functional heterogeneity within the human SN (101,103,106), revealing a complex network whereby the dopaminergic system, through distinct subregions of the SN, navigates the confluence of various types of salience to modulate behaviour and memory processes. Specifically, the dorsal SN predominantly projects to striatal areas, which in turn modulate executive and attentional functions, while the ventral SN extends projections to the hippocampus and amygdala, which are crucial for encoding salient events into memory (106). This distinction aligns with our observation of the dorsal SN’s involvement in processing salience related to reward or unexpectedness, and prior studies showing its role in visuo-motor-related learning (101). On the other hand, the strong connectivity of the ventral SN to cortical areas such as the caudate, cingulate, and insula (101,106) in addition to hippocampus and amygdala might in turn explain its role in mediating the effects of unexpectedness on memory outcomes.

In summary, our behavioural results suggest distinct effects of reward- and unexpectedness-related salience, manifesting respectively as response biases and enhanced memory. At the same time, we were able to identify distinct brain networks associated with different types of salience, as well as networks involved in processing salience and modulating memory encoding. Reward- and unexpectedness-related brain networks largely overlapped with the expected reward and salience networks (cf. Figure 5, Supplementary Tables 4 and 5, Supplementary Results 3). An interesting distinction was observed within the cingulate cortex: The posterior regions were predominantly involved in unexpected-related salience, while the anterior regions engaged in both reward- and unexpectedness-related salience. Although the expected distinction between the SN and locus coeruleus in supporting reward and contextual unexpectedness, respectively, could not be verified in this study, we confirmed the functional implications of anatomical subregions within the SN. Processing stimulus salience, regardless of the type, preferentially engaged the dorsal SN, while salience-associated memory encoding appeared to be more supported by the ventral SN.

### 4.3. Limitations and Considerations for Future Research

This study is not without its limitations. Given the 100% reward allocation with the reward-associated category, our reward manipulation was likely to have been predictable, which could have tempered our reward-associated salience effect by reducing the influence of prediction errors. Rouhani et al.’s work provides an intricate understanding of this dynamic; they found that cues associated with higher RPEs at the moment of cue presentation were better remembered as learning progressed (122). In their experiment, they were able to dissociate the effects of cue values and RPEs on memory, establishing that an RPE signal is essential for the mnemonic enhancement of cue events (122). As our study’s intention was to disentangle the neural correlates of two salience types, a deterministic association between the reward and its respective category was necessary to create a reward anticipation effect that could be contrasted with the inherently unpredictable nature of contextually unexpected events. This affected our ability to investigate RPE-dependent effects. Future studies focusing on midbrain and brainstem function should systematically alter stimulus and reward expectedness in order to compare reward, prediction error and frequency effects.

Lastly, given our aim to compare two different types of salience associated with dopaminergic and noradrenergic modulation, reward and contextual unexpectedness, our task necessarily resulted in differential behavioural correlates of salience. While infrequently presented stimuli, in line with von Restorff effect (26–28,44–46), primarily elicited an enhanced memory effect, reward associations reward associations predominantly affected response biases. This made a comparison of the extent of salience manipulations difficult, limiting us to a qualitative comparison. Nonetheless, even in the absence of comparable behavioural memory effects, activity patterns for successfully encoded scenes across reward-associated and infrequently presented scenes significantly overlapped (Jaccard Index = 0.5807; overlapping activations indicated by white outlines in Supplementary Figure 5). This suggests that comparable networks for memory encoding across salience types might be recruited. Simultaneously, whether similar response bias effects could be observed in relation to contextually unexpected stimuli remains questionable, as response bias modulation appears to be more specifically linked to reward associations (57). Nevertheless, future studies should also aim to allow for a comparison of more quantitative aspects of different types of salience and their effects on brainstem or midbrain function. This could, for example, be achieved by including additional measures of arousal, such as pupillometry or skin conductance charges, if behavioural correlates cannot be equated.

## 5 Conclusion

In conclusion, our study delineates both unique and overlapping networks involved in the processing and memory encoding of contextual unexpectedness-related and reward-related salience. Utilising an MRI analysis pipeline optimised for enhanced spatial precision in assessing the neuromodulatory structures in the midbrain and brainstem, we observed differential engagement of regions traditionally associated with dopaminergic modulation in processing distinct types of salience. Future studies, perhaps focusing on probabilistic reward schemes or a wider array of events such as negative or shocking incidents, can further consolidate our understanding of not only neuromodulatory structures’ differential involvement but also their interactive roles in modulating responses to salient events.

## References

1. Aston-Jones G, Cohen JD. Adaptive gain and the role of the locus coeruleus-norepinephrine system in optimal performance. J Comp Neurol. 2005 Dec 5;493(1):99–110.

2. Sara SJ. The locus coeruleus and noradrenergic modulation of cognition. Nat Rev Neurosci. 2009 Mar;10(3):211–23.

3. Schultz W. Behavioral dopamine signals. Trends Neurosci. 2007 May;30(5):203– 10.

4. Berridge CW, Waterhouse BD. The locus coeruleus–noradrenergic system: modulation of behavioral state and state-dependent cognitive processes. Brain Res Rev. 2003 Apr;42(1):33–84.

5. Robbins TW, Arnsten AFT. The Neuropsychopharmacology of Fronto-Executive Function: Monoaminergic Modulation. Annu Rev Neurosci. 2009 Jun 1;32(1):267–87.

6. Schultz W. Neuronal Reward and Decision Signals: From Theories to Data. Physiol Rev. 2015 Jul;95(3):853–951.

7. Blier P, El Mansari M. Serotonin and beyond: therapeutics for major depression. Philos Trans R Soc B Biol Sci. 2013 Apr 5;368(1615):20120536.

8. Arnsten AFT. Catecholamine Influences on Dorsolateral Prefrontal Cortical Networks. Biol Psychiatry. 2011 Jun;69(12):e89–99.

9. Berridge CW, Schmeichel BE, España RA. Noradrenergic modulation of wakefulness/arousal. Sleep Med Rev. 2012 Apr;16(2):187–97.

10. Düzel E, Bunzeck N, Guitart-Masip M, Düzel S. NOvelty-related Motivation of Anticipation and exploration by Dopamine (NOMAD): Implications for healthy aging. Neurosci Biobehav Rev. 2010 Apr;34(5):660–9.

11. Hämmerer D, Callaghan MF, Hopkins A, Kosciessa J, Betts M, Cardenas-Blanco A, et al. Locus coeruleus integrity in old age is selectively related to memories linked with salient negative events. Proc Natl Acad Sci. 2018 Feb 27;115(9):2228–33.

12. Lisman JE, Grace AA. The Hippocampal-VTA Loop: Controlling the Entry of Information into Long-Term Memory. Neuron. 2005 Jun;46(5):703–13.

13. Luo AH, Tahsili-Fahadan P, Wise RA, Lupica CR, Aston-Jones G. Linking Context with Reward: A Functional Circuit from Hippocampal CA3 to Ventral Tegmental Area. Science. 2011 Jul 15;333(6040):353–7.

14. Samson Y, Wu JJ, Friedman AH, Davis JN. Catecholaminergic innervation of the hippocampus in the cynomolgus monkey. J Comp Neurol. 1990 Aug 8;298(2):250–63.

15. Schott BH, Sellner DB, Lauer CJ, Habib R, Frey JU, Guderian S, et al. Activation of Midbrain Structures by Associative Novelty and the Formation of Explicit Memory in Humans. Learn Mem. 2004 Jul;11(4):383–7.

16. Shohamy D, Adcock RA. Dopamine and adaptive memory. Trends Cogn Sci. 2010 Oct;14(10):464–72.

17. Aston-Jones G, Cohen JD. AN INTEGRATIVE THEORY OF LOCUS COERULEUS-NOREPINEPHRINE FUNCTION: Adaptive Gain and Optimal Performance. Annu Rev Neurosci. 2005 Jul 21;28(1):403–50.

18. Doya K. Modulators of decision making. Nat Neurosci. 2008 Apr;11(4):410–6.

19. Grace AA. Dysregulation of the dopamine system in the pathophysiology of schizophrenia and depression. Nat Rev Neurosci. 2016 Aug;17(8):524–32.

20. McDevitt RA, Tiran-Cappello A, Shen H, Balderas I, Britt JP, Marino RAM, et al. Serotonergic versus Nonserotonergic Dorsal Raphe Projection Neurons: Differential Participation in Reward Circuitry. Cell Rep. 2014 Sep;8(6):1857–69.

21. Schomaker J, Meeter M. Short- and long-lasting consequences of novelty, deviance and surprise on brain and cognition. Neurosci Biobehav Rev. 2015 Aug;55:268–79.

22. Takeuchi T, Duszkiewicz AJ, Sonneborn A, Spooner PA, Yamasaki M, Watanabe M, et al. Locus coeruleus and dopaminergic consolidation of everyday memory. Nature. 2016 Sep;537(7620):357–62.

23. O’Carroll CM, Martin SJ, Sandin J, Frenguelli B, Morris RGM. Dopaminergic modulation of the persistence of one-trial hippocampus-dependent memory. Learn Mem. 2006 Nov;13(6):760–9.

24. Froemke RC. Plasticity of Cortical Excitatory-Inhibitory Balance. Annu Rev Neurosci. 2015 Jul 8;38(1):195–219.

25. Lisman J, Grace AA, Duzel E. A neoHebbian framework for episodic memory; role of dopamine-dependent late LTP. Trends Neurosci. 2011 Oct;34(10):536–47.

26. Duszkiewicz AJ, McNamara CG, Takeuchi T, Genzel L. Novelty and Dopaminergic Modulation of Memory Persistence: A Tale of Two Systems. Trends Neurosci. 2019 Feb;42(2):102–14.

27. Yamasaki M, Takeuchi T. Locus Coeruleus and Dopamine-Dependent Memory Consolidation. Neural Plast. 2017;2017:1–15.

28. Devoto P, Flore G, Saba P, Fa M, Gessa GL. Stimulation of the locus coeruleus elicits noradrenaline and dopamine release in the medial prefrontal and parietal cortex. J Neurochem. 2005 Jan;92(2):368–74.

29. Adcock RA, Thangavel A, Whitfield-Gabrieli S, Knutson B, Gabrieli JDE. Reward-Motivated Learning: Mesolimbic Activation Precedes Memory Formation. Neuron. 2006 May;50(3):507–17.

30. Barto A, Mirolli M, Baldassarre G. Novelty or Surprise? Front Psychol [Internet]. 2013 [cited 2023 Feb 7];4. Available from: http://journal.frontiersin.org/article/10.3389/fpsyg.2013.00907/abstract

31. Bunzeck N, Düzel E. Absolute Coding of Stimulus Novelty in the Human Substantia Nigra/VTA. Neuron. 2006 Aug;51(3):369–79.

32. Ikemoto S. Dopamine reward circuitry: Two projection systems from the ventral midbrain to the nucleus accumbens–olfactory tubercle complex. Brain Res Rev. 2007 Nov;56(1):27–78.

33. Kafkas A, Montaldi D. Striatal and midbrain connectivity with the hippocampus selectively boosts memory for contextual novelty. Hippocampus. 2015 Nov;25(11):1262–73.

34. Wittmann BC, Bunzeck N, Dolan RJ, Düzel E. Anticipation of novelty recruits reward system and hippocampus while promoting recollection. NeuroImage. 2007 Oct;38(1):194–202.

35. Liu KY, Marijatta F, Hämmerer D, Acosta-Cabronero J, Düzel E, Howard RJ. Magnetic resonance imaging of the human locus coeruleus: A systematic review. Neurosci Biobehav Rev. 2017 Dec;83:325–55.

36. Yi YJ, Lüsebrink F, Ludwig M, Maaß A, Ziegler G, Yakupov R, et al. It is the locus coeruleus! Or… is it?: a proposition for analyses and reporting standards for structural and functional magnetic resonance imaging of the noradrenergic locus coeruleus. Neurobiol Aging. 2023 Sep;129:137–48.

37. Khosla A, Raju AS, Torralba A, Oliva A. Understanding and Predicting Image Memorability at a Large Scale. In: 2015 IEEE International Conference on Computer Vision (ICCV) [Internet]. Santiago, Chile: IEEE; 2015 [cited 2023 Jun 21]. p. 2390–8. Available from: http://ieeexplore.ieee.org/document/7410632/

38. Fonov V, Evans AC, Botteron K, Almli CR, McKinstry RC, Collins DL. Unbiased average age-appropriate atlases for pediatric studies. NeuroImage. 2011 Jan;54(1):313–27.

39. Behzadi Y, Restom K, Liau J, Liu TT. A component based noise correction method (CompCor) for BOLD and perfusion based fMRI. NeuroImage. 2007 Aug;37(1):90–101.

40. Genovese CR, Lazar NA, Nichols T. Thresholding of Statistical Maps in Functional Neuroimaging Using the False Discovery Rate. NeuroImage. 2002 Apr;15(4):870–8.

41. Beissner F, Baudrexel S. Investigating the human brainstem with structural and functional MRI. Front Hum Neurosci. 2014;

42. Pauli WM, Nili AN, Tyszka JM. A high-resolution probabilistic in vivo atlas of human subcortical brain nuclei. Sci Data. 2018 Apr 17;5(1):180063.

43. Hautus MJ. Corrections for extreme proportions and their biasing effects on estimated values ofd’. Behav Res Methods Instrum Comput. 1995 Mar;27(1):46– 51.

44. Kafkas A, Montaldi D. How do memory systems detect and respond to novelty? Neurosci Lett. 2018 Jul;680:60–8.

45. Levy BJ, Wagner AD. Cognitive control and right ventrolateral prefrontal cortex: reflexive reorienting, motor inhibition, and action updating: Cognitive control and right ventrolateral PFC. Ann N Y Acad Sci. 2011 Apr;1224(1):40–62.

46. von Restorff H. Über die Wirkung von Bereichsbildungen im Spurenfeld. Psychol Forsch. 1933;18:299–342.

47. Verschuere B, Kleinberg B, Theocharidou K. RT-based memory detection: Item saliency effects in the single-probe and the multiple-probe protocol. J Appl Res Mem Cogn. 2015 Mar;4(1):59–65.

48. Kirchhoff BA, Wagner AD, Maril A, Stern CE. Prefrontal–Temporal Circuitry for Episodic Encoding and Subsequent Memory. J Neurosci. 2000 Aug 15;20(16):6173–80.

49. Kormi-Nouri R, Nilsson LG, Ohta N. The BlackwellPublishing,Ltd. novelty effect: Support for the Novelty-Encoding Hypothesis. Scand J Psychol. 2005;

50. Ranganath C, Rainer G. Neural mechanisms for detecting and remembering novel events. Nat Rev Neurosci. 2003 Mar;4(3):193–202.

51. Gimbel SI, Brewer JB. Reaction time, memory strength, and fMRI activity during memory retrieval: Hippocampus and default network are differentially responsive during recollection and familiarity judgments. Cogn Neurosci. 2011 Jan 27;2(1):19–26.

52. Dewhurst SA, Holmes SJ, Brandt KR, Dean GM. Measuring the speed of the conscious components of recognition memory: Remembering is faster than knowing. Conscious Cogn. 2006 Mar;15(1):147–62.

53. Gimbel SI, Brewer JB. Reaction time, memory strength, and fMRI activity during memory retrieval: Hippocampus and default network are differentially responsive during recollection and familiarity judgments. Cogn Neurosci. 2011 Jan 27;2(1):19–26.

54. Rouhani N, Norman KA, Niv Y. Dissociable effects of surprising rewards on learning and memory. J Exp Psychol Learn Mem Cogn. 2018 Sep;44(9):1430– 43.

55. Schultz W, Dayan P, Montague PR. A Neural Substrate of Prediction and Reward. Science. 1997 Mar 14;275(5306):1593–9.

56. Wimmer GE, Braun EK, Daw ND, Shohamy D. Episodic Memory Encoding Interferes with Reward Learning and Decreases Striatal Prediction Errors. J Neurosci. 2014 Nov 5;34(45):14901–12.

57. Bowen HJ, Marchesi ML, Kensinger EA. Reward motivation influences response bias on a recognition memory task. Cognition. 2020 Oct;203:104337.

58. Beierholm U, Guitart-Masip M, Economides M, Chowdhury R, Düzel E, Dolan R, et al. Dopamine Modulates Reward-Related Vigor. Neuropsychopharmacology. 2013 Jul;38(8):1495–503.

59. Guitart-Masip M, Fuentemilla L, Bach DR, Huys QJM, Dayan P, Dolan RJ, et al. Action Dominates Valence in Anticipatory Representations in the Human Striatum and Dopaminergic Midbrain. J Neurosci. 2011 May 25;31(21):7867–75.

60. Atkinson RC, Shiffrin RM. HUMAN MEMORY: A PROPOSED SYSTEM AND ITS CONTROL PROCESSES. In: Human Memory [Internet]. Elsevier; 1968 [cited 2023 Jun 22]. p. 7–113. Available from: https://linkinghub.elsevier.com/retrieve/pii/B9780121210502500065

61. Staudigl T, Hanslmayr S. Theta Oscillations at Encoding Mediate the Context-Dependent Nature of Human Episodic Memory. Curr Biol. 2013 Jun;23(12):1101– 6.

62. Tulving E, Thomson DM. Encoding specificity and retrieval processes in episodic memory. Psychol Rev. 1973 Sep;80(5):352–73.

63. Wittmann BC, Schott BH, Guderian S, Frey JU, Heinze HJ, Düzel E. Reward-Related fMRI Activation of Dopaminergic Midbrain Is Associated with Enhanced Hippocampus-Dependent Long-Term Memory Formation. Neuron. 2005 Feb;45(3):459–67.

64. Guitart-Masip M, Bunzeck N, Stephan KE, Dolan RJ, Duzel E. Contextual Novelty Changes Reward Representations in the Striatum. J Neurosci. 2010 Feb 3;30(5):1721–6.

65. Kiehl KA, Laurens KR, Duty TL, Forster BB, Liddle PF. An Event-Related fMRI Study of Visual and Auditory Oddball Tasks. J Psychophysiol. 2001 Oct;15(4):221–40.

66. Seeley WW, Menon V, Schatzberg AF, Keller J, Glover GH, Kenna H, et al. Dissociable Intrinsic Connectivity Networks for Salience Processing and Executive Control. J Neurosci. 2007 Feb 28;27(9):2349–56.

67. Hollerman JR, Tremblay L, Schultz W. Involvement of basal ganglia and orbitofrontal cortex in goal-directed behavior. In: Progress in Brain Research [Internet]. Elsevier; 2000 [cited 2023 Feb 3]. p. 193–215. Available from: https://linkinghub.elsevier.com/retrieve/pii/S0079612300260159

68. Liu X, Hairston J, Schrier M, Fan J. Common and distinct networks underlying reward valence and processing stages: A meta-analysis of functional neuroimaging studies. Neurosci Biobehav Rev. 2011 Apr;35(5):1219–36.

69. Samanez-Larkin GR, Gibbs SEB, Khanna K, Nielsen L, Carstensen LL, Knutson B. Anticipation of monetary gain but not loss in healthy older adults. Nat Neurosci. 2007 Jun;10(6):787–91.

70. Rolls ET. The Orbitofrontal Cortex and Reward. Cereb Cortex. 2000 Mar 1;10(3):284–94.

71. Krebs RM, Park HRP, Bombeke K, Boehler CN. Modulation of locus coeruleus activity by novel oddball stimuli. Brain Imaging Behav. 2018 Apr;12(2):577–84.

72. Sara SJ, Vankov A, Hervé A. Locus Coeruleus-evoked Responses in Behaving Rats: A Clue to the Role of Noradrenaline in Memory. Brain Res Bull. 1994;35:457–65.

73. Rigoli F, Friston KJ, Dolan RJ. Neural processes mediating contextual influences on human choice behaviour. Nat Commun. 2016 Aug 18;7(1):12416.

74. Daffner KR, Mesulam MM, Scinto LFM, Acar D, Calvo V, Faust R, et al. The central role of the prefrontal cortex in directing attention to novel events. Brain. 2000 May;123(5):927–39.

75. Hawco C, Lepage M. Overlapping patterns of neural activity for different forms of novelty in fMRI. Front Hum Neurosci [Internet]. 2014 Sep 8 [cited 2023 Feb 3];8. Available from: http://journal.frontiersin.org/article/10.3389/fnhum.2014.00699/abstract

76. Kirino E, Belger A, Goldman-Rakic P, McCarthy G. Prefrontal Activation Evoked by Infrequent Target and Novel Stimuli in a Visual Target Detection Task: An Event-Related Functional Magnetic Resonance Imaging Study. J Neurosci. 2000 Sep 1;20(17):6612–8.

77. Corbetta M, Shulman GL. Control of goal-directed and stimulus-driven attention in the brain. Nat Rev Neurosci. 2002 Mar 1;3(3):201–15.

78. Gogolla N. The insular cortex. Curr Biol. 2017 Jun;27(12):R580–6.

79. Kafkas A, Montaldi D. Two separate, but interacting, neural systems for familiarity and novelty detection: A dual-route mechanism: Familiarity and Novelty Detection Processes. Hippocampus. 2014 May;24(5):516–27.

80. Shuman M, Kanwisher N. Numerical Magnitude in the Human Parietal Lobe: Tests of Representational Generality and Domain Specificity. Neuron. 2004 Oct 28;44(557–569).

81. Uddin LQ. Salience processing and insular cortical function and dysfunction. Nat Rev Neurosci. 2015 Jan;16(1):55–61.

82. Haber SN. The primate basal ganglia: parallel and integrative networks. J Chem Neuroanat. 2003 Dec;26(4):317–30.

83. Kamiński J, Mamelak AN, Birch K, Mosher CP, Tagliati M, Rutishauser U. Novelty-Sensitive Dopaminergic Neurons in the Human Substantia Nigra Predict Success of Declarative Memory Formation. Curr Biol. 2018 May;28(9):1333–1343.e4.

84. Mikell CB, Sheehy JP, Youngerman BE, McGovern RA, Wojtasiewicz TJ, Chan AK, et al. Features and timing of the response of single neurons to novelty in the substantia nigra. Brain Res. 2014 Jan;1542:79–84.

85. D’Esposito M, Postle BR, Ballard D, Lease J. Maintenance versus Manipulation of Information Held in Working Memory: An Event-Related fMRI Study. Brain Cogn. 1999 Oct;41(1):66–86.

86. Jastorff J, Clavagnier S, Gergely G, Orban GA. Neural Mechanisms of Understanding Rational Actions: Middle Temporal Gyrus Activation by Contextual Violation. Cereb Cortex. 2011 Feb;21(2):318–29.

87. Paulus MP, Feinstein JS, Castillo G, Simmons AN, Stein MB. Dose-Dependent Decrease of Activation in Bilateral Amygdala and Insula by Lorazepam During Emotion Processing. Arch Gen Psychiatry. 2005 Mar 1;62(3):282.

88. Bush G, Vogt BA, Holmes J, Dale AM, Greve D, Jenike MA, et al. Dorsal anterior cingulate cortex: A role in reward-based decision making. Proc Natl Acad Sci. 2002 Jan 8;99(1):523–8.

89. Domic-Siede M, Irani M, Valdés J, Perrone-Bertolotti M, Ossandón T. Theta activity from frontopolar cortex, mid-cingulate cortex and anterior cingulate cortex shows different roles in cognitive planning performance. NeuroImage. 2021 Feb;226:117557.

90. Shenhav A, Cohen JD, Botvinick MM. Dorsal anterior cingulate cortex and the value of control. Nat Neurosci. 2016 Oct;19(10):1286–91.

91. Jang AI, Nassar MR, Dillon DG, Frank MJ. Positive reward prediction errors during decision-making strengthen memory encoding. Nat Hum Behav. 2019 May 6;3(7):719–32.

92. Stanek JK, Dickerson KC, Chiew KS, Clement NJ, Adcock RA. Expected Reward Value and Reward Uncertainty Have Temporally Dissociable Effects on Memory Formation. J Cogn Neurosci. 2019 Oct;31(10):1443–54.

93. Klein I, Paradis AL, Poline JB, Kosslyn SM, Le Bihan D. Transient Activity in the Human Calcarine Cortex During Visual-Mental Imagery: An Event-Related fMRI Study. J Cogn Neurosci. 2000 Nov 1;12(Supplement 2):15–23.

94. Wojciulik E, Kanwisher N. The Generality of Parietal Involvement in Visual Attention. Neuron. 1999 Aug;23(4):747–64.

95. Cabeza R, Ciaramelli E, Olson IR, Moscovitch M. The parietal cortex and episodic memory: an attentional account. Nat Rev Neurosci. 2008 Aug;9(8):613–25.

96. Blumenfeld RS, Ranganath C. Prefrontal Cortex and Long-Term Memory Encoding: An Integrative Review of Findings from Neuropsychology and Neuroimaging. The Neuroscientist. 2007 Jun;13(3):280–91.

97. Grill-Spector K, Weiner KS. The functional architecture of the ventral temporal cortex and its role in categorization. Nat Rev Neurosci. 2014 Aug;15(8):536–48.

98. Hoffman KL, Logothetis NK. Cortical mechanisms of sensory learning and object recognition. Philos Trans R Soc B Biol Sci. 2009 Feb 12;364(1515):321–9.

99. Meyer T, Olson CR. Statistical learning of visual transitions in monkey inferotemporal cortex. Proc Natl Acad Sci. 2011 Nov 29;108(48):19401–6.

100. Kringelbach M, Rolls ET. The functional neuroanatomy of the human orbitofrontal cortex: evidence from neuroimaging and neuropsychology. Prog Neurobiol. 2004 Apr;72(5):341–72.

101. Zhang Y, Larcher KMH, Misic B, Dagher A. Anatomical and functional organization of the human substantia nigra and its connections. eLife. 2017 Aug 21;6:e26653.

102. Krebs RM, Heipertz D, Schuetze H, Duzel E. Novelty increases the mesolimbic functional connectivity of the substantia nigra/ventral tegmental area (SN/VTA) during reward anticipation: Evidence from high-resolution fMRI. NeuroImage. 2011 Sep;58(2):647–55.

103. Wittmann BC, Schott BH, Guderian S, Frey JU, Heinze HJ, Düzel E. Reward-Related fMRI Activation of Dopaminergic Midbrain Is Associated with Enhanced Hippocampus-Dependent Long-Term Memory Formation. Neuron. 2005 Feb;45(3):459–67.

104. Adcock RA, Thangavel A, Whitfield-Gabrieli S, Knutson B, Gabrieli JDE. Reward-Motivated Learning: Mesolimbic Activation Precedes Memory Formation. Neuron. 2006 May;50(3):507–17.

105. Hämmerer D, Schwartenbeck P, Gallagher M, FitzGerald THB, Düzel E, Dolan RJ. Older adults fail to form stable task representations during model-based reversal inference. Neurobiol Aging. 2019 Feb;74:90–100.

106. Haber SN, Knutson B. The Reward Circuit: Linking Primate Anatomy and Human Imaging. Neuropsychopharmacology. 2010 Jan;35(1):4–26.

107. O’Doherty JP. Reward representations and reward-related learning in the human brain: insights from neuroimaging. Curr Opin Neurobiol. 2004 Dec;14(6):769–76.

108. Grahn JA, Parkinson JA, Owen AM. The cognitive functions of the caudate nucleus. Prog Neurobiol. 2008 Nov;86(3):141–55.

109. Graybiel AM. Habits, Rituals, and the Evaluative Brain. Annu Rev Neurosci. 2008 Jul 1;31(1):359–87.

110. Yanike M, Ferrera VP. Representation of Outcome Risk and Action in the Anterior Caudate Nucleus. J Neurosci. 2014 Feb 26;34(9):3279–90.

111. Williams ZM, Eskandar EN. Selective enhancement of associative learning by microstimulation of the anterior caudate. Nat Neurosci. 2006 Apr;9(4):562–8.

112. Pihlajamaki M, Tanila H, Kononen M, Hanninen T, Hamalainen A, Soininen H, et al. Visual presentation of novel objects and new spatial arrangements of objects differentially activates the medial temporal lobe subareas in humans. Eur J Neurosci. 2004 Apr;19(7):1939–49.

113. Kaplan R, Horner AJ, Bandettini PA, Doeller CF, Burgess N. Human hippocampal processing of environmental novelty during spatial navigation: Human Hippocampal Processing Of Environmental Novelty. Hippocampus. 2014 Jul;24(7):740–50.

114. Aminoff E, Gronau N, Bar M. The Parahippocampal Cortex Mediates Spatial and Nonspatial Associations. Cereb Cortex. 2007 Jul 1;17(7):1493–503.

115. Baumann O, Mattingley JB. Functional Organization of the Parahippocampal Cortex: Dissociable Roles for Context Representations and the Perception of Visual Scenes. J Neurosci. 2016 Feb 24;36(8):2536–42.

116. Takeuchi T, Duszkiewicz AJ, Sonneborn A, Spooner PA, Yamasaki M, Watanabe M, et al. Locus coeruleus and dopaminergic consolidation of everyday memory. Nature. 2016 Sep;537(7620):357–62.

117. Ludwig M, Hammerer D, Lüsebrink F, Düzel E. Interrogating the role of the noradrenergic locus coeruleus in memory encoding in aging: Neuroimaging / Optimal neuroimaging measures for early detection. Alzheimers Dement. 2020 Dec;16(S5):e044039.

118. Hampson M, Driesen NR, Skudlarski P, Gore JC, Constable RT. Brain Connectivity Related to Working Memory Performance. J Neurosci. 2006 Dec 20;26(51):13338–43.

119. McCoy AN, Crowley JC, Haghighian G, Dean HL, Platt ML. Saccade Reward Signals in Posterior Cingulate Cortex. Neuron. 2003 Dec;40(5):1031–40.

120. Menon V, Uddin LQ. Saliency, switching, attention and control: a network model of insula function. Brain Struct Funct. 2010 Jun;214(5–6):655–67.

121. Sridharan D, Levitin DJ, Menon V. A critical role for the right fronto-insular cortex in switching between central-executive and default-mode networks. Proc Natl Acad Sci. 2008 Aug 26;105(34):12569–74.

122. Rouhani N, Niv Y. Signed and unsigned reward prediction errors dynamically enhance learning and memory. eLife. 2021 Mar 4;10:e61077.

