## Supplementary Materials for "Decoding Salience: A Functional Magnetic Resonance Imaging Investigation of Reward and Contextual Unexpectedness in Memory Encoding and Retrieval"

Yeo-Jin Yi[ab*], Michael C. Kreißl[bc], Oliver Speck[bdef], Emrah Düzel[†aeg], Dorothea Hämmerer[†aegh] ([†]shared last)

[a] Institute of Cognitive Neurology and Dementia Research, Otto-von-Guericke University, Magdeburg, Germany
[b] German Center for Neurodegenerative Diseases (DZNE), Magdeburg, Germany
[c] Division of Nuclear Medicine, Department of Nuclear Medicine, Otto-von-Guericke University, Magdeburg, Germany
[d] Biomedical Magnetic Resonance, Faculty of Natural Sciences, Otto-von-Guericke University, Magdeburg, Germany
[e] Center for Behavioral Brain Sciences, Magdeburg, Germany
[f] Leibniz Institute for Neurobiology, Magdeburg, Germany
[g] Institute of Cognitive Neuroscience, University College London, United Kingdom
[h] Department of Psychology, University of Innsbruck, Innsbruck, Austria

***Correspondence:**
Yeo-Jin Yi


**Keywords: fMRI, Reward, Contextual Unexpectedness, Memory, Midbrain, Cognition**

### Abstract

The present study investigated the neuromodulatory substrates of salience processing and its impact on memory encoding and behaviour, with a specific focus on two distinct types of salience: reward and contextual unexpectedness. 46 participants performed a novel task paradigm modulating these two aspects independently and allowing for investigating their distinct and interactive effects on memory encoding while undergoing high resolution fMRI. By using advanced image processing techniques tailored to examine midbrain and brainstem nuclei with high precision, our study additionally aimed to elucidate differential activation patterns in subcortical nuclei in response to reward-associated and contextually unexpected stimuli, including distinct pathways involving in particular dopaminergic modulation. We observed a differential involvement of the ventral striatum, substantia nigra and caudate nucleus, as well as a functional specialisation within the subregions of the cingulate cortex for the two salience types. Moreover, distinct subregions within the substantia nigra in processing salience could be identified. Dorsal areas preferentially processed salience related to stimulus processing (of both reward and contextual unexpectedness) versus ventral areas were involved in salience-related memory encoding (for contextual unexpectedness only). These functional specialisations within SN are in line with different projection patterns of dorsal and ventral SN to brain areas supporting attention and memory, respectively. By disentangling stimulus processing and memory encoding related to two salience types, we hope to further consolidate our understanding of neuromodulatory structures' differential as well as interactive roles in modulating behavioural responses to salient events.

### 1. Supplementary Method 1: Overview of the spatial transformation procedure and parameters

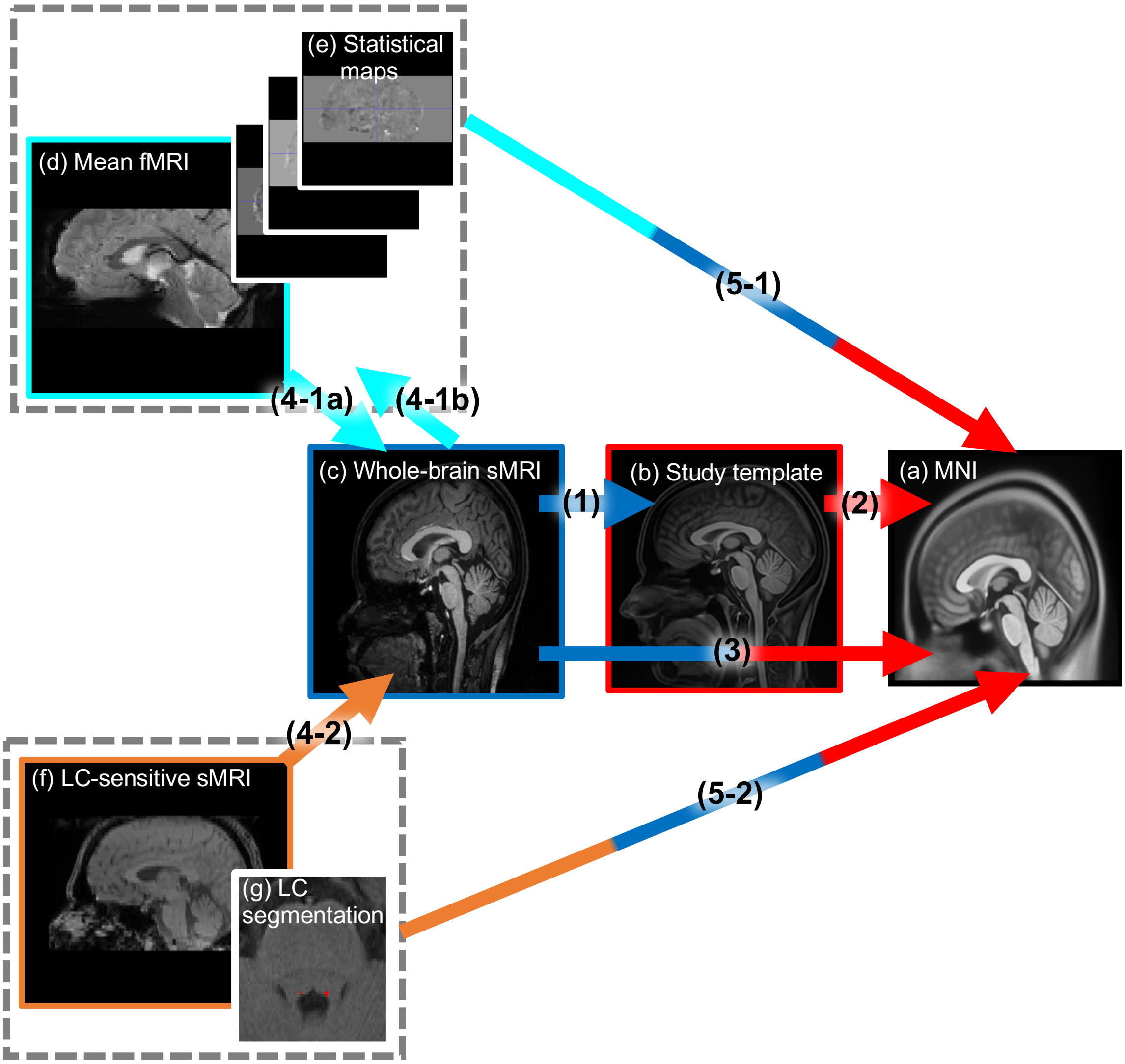


**Supplementary Figure 1. Overview of the spatial transformation steps. Adapted from Yi et al. (1).**

Single subject data in native space (c-g) are transformed into Montreal Neurological Institute (MNI) structural template space (a) for group-level analyses. Specifically, (a) represents MNI space as created by Fonov and colleagues (2). (b) refers to a study-specific template space, which is generated from all whole-brain structural images in the dataset that are corrected for RF-field-related inhomogeneity with *N4BiasFieldCorrection* function of Advanced Normalization Tools (parameters:

[image dimension=3, shrink factor=2, convergence iterations=200x150x100x50, convergence threshold=$10^{-6}$, spline spacing=200]; ANTs; (3), using *antsMultivariateTemplateConstruction2.sh* function also from ANTs (parameters: image dimension=3, number of modalities=1, rigid-body registration on, smoothing factors=4x2x1x0, the rest of the parameters were kept at the default value). The study-specific template assists in the transformation from native to MNI space as an intermediate step. (c) is a whole-brain structural image, while (d) denotes the mean functional image post-reslicing, realignment, and unwarping. (e) represents statistical maps generated from smoothed functional images after applying general linear models (GLMs) using Statistical Parametric Mapping 12 (SPM12; http://www.fil.ion.ucl.ac.uk/spm12.html). (f) is a neuromelanin-sensitive structural image used for locus coeruleus (LC) imaging, and (g) depicts a manually segmented LC mask drawn in red on image (f). Arrows indicate spatial transformation steps, with arrow heads pointing towards the image space into which an image is transformed. Arrows featuring multiple colours indicate concatenated transformation matrices; for example, the green-blue-magenta arrow (5-1) demonstrates the combination of transformation matrices calculated from the mean functional image (d) to the structural image (c) or vice versa (steps 4-1a and 4-1b), the structural image (c) to study-specific template (b) and MNI (step 1), and the study-specific template (b) to MNI (a) (step 2) into one transformation step (step 5-1). This process transforms the statistical maps (e) and the mean functional image (d) into MNI space (a) in one single step. The numbers represent the order of transformation steps executed in the pipeline. Steps 4-1a and 4-1b correspond to the recommended transformation alternatives for whole-brain and partial volume functional data, respectively. The scripts used for this processing pipeline are available for download from this Github repository: https://github.com/alex-yi-writes/LC-SpatialTransformation2021

### 2. Supplementary Method 2: Functional image quality assessment and quantification procedure

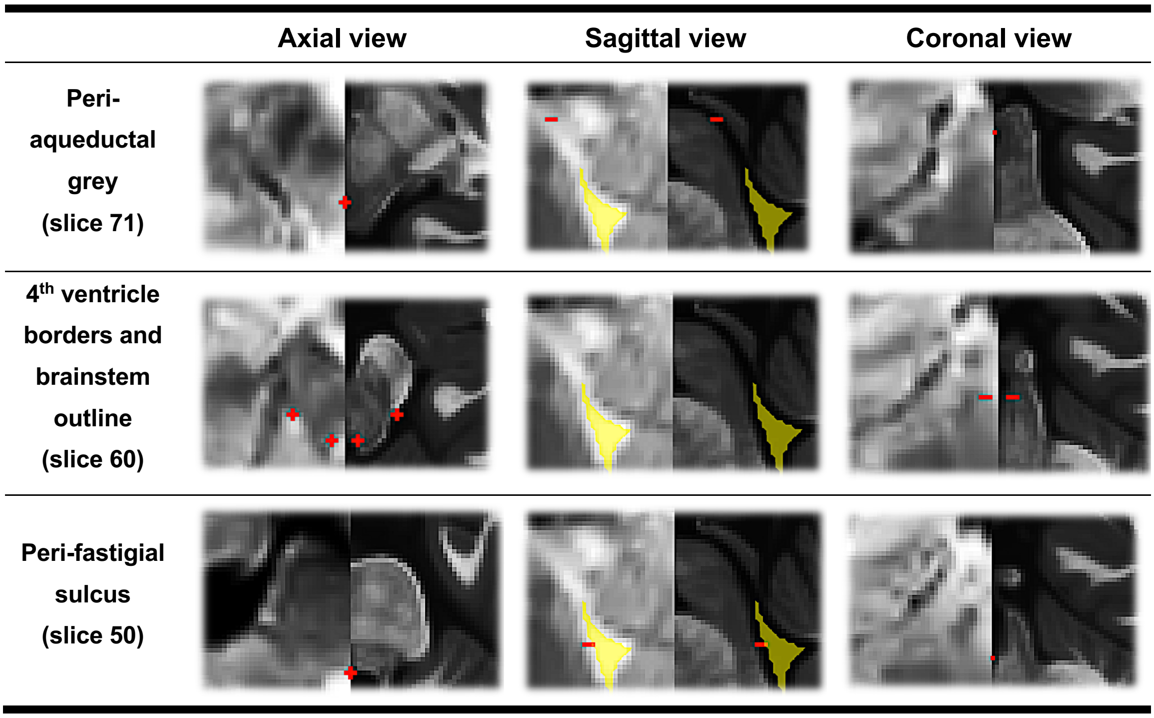


**Supplementary Figure 2. Landmark placement for quality control of functional data. Adapted from Yi et al. (1).**

In order to assess the precision of the transformations applied to functional images (Suppl. Figure 1d) and statistical maps (Suppl. Figure 1e), it was necessary to establish anatomical landmarks within the structural MNI template space. These landmarks should be relevant to the brainstem and midbrain areas and must be clearly visible in the normalised functional images, as the neuromodulatory structures themselves are not directly visible. Such landmarks may encompass clearly discernible structures relevant to our study's neuromodulatory nuclei of interest. For instance, the border between the cerebrospinal fluid and the brainstem of the fourth ventricle is significant for anatomically locating the locus coeruleus (see Suppl. Figure 2, second row). Similarly, the periaqueductal grey is crucial in aligning images to the substantia nigra and the ventral tegmental area. These chosen landmarks, saved as segmentations, could then be individually compared to pre-set landmarks on the MNI space (Suppl. Figure 1a).

After the assessment of the entire dataset is completed, all the landmarks aggregated in the same space was visually inspected by generating a heatmap of the average of the saved individual landmarks. In addition, a frame-by-frame video of the transformed images was created, which facilitated easy identification of outliers. Any registration or normalisation errors identified at this step were rectified individually, depending on the nature of each error. For instance, the brain in the transformed mean functional image may extend beyond the tissue area of the MNI template following spatial transformation. By retracing the transformation steps, the error source was identified during step 4-1 (as shown in Suppl. Figure 1), which is the rigid registration of the structural whole-brain image to the mean functional image. This issue appears to occur due to the *antsRegistrationSyN.sh* function utilised in this step, as it interprets the intensity of the tissues near the skull in the mean functional image as more similar to the structural image's skull than the faint skull traces in the mean functional image. This problem could be resolved by skull-stripping the images involved in the registration process.

By aggregating the in-plane distances between the pre-set landmarks and each subject’s landmarks using a custom MATLAB script, we could quantify and report the quality of spatial transformations in group space in each landmark (see Figure 3 of the main text). The codes utilised for this procedure are available for download from this Github repository: https://github.com/alex-yi-writes/LC-SpatialTransformation2021

However, it is important to note that any deviations observed at this stage of spatial transformations could result from a range of factors, including a) low native image resolution, which may affect the precision of transformations and quality assessments, b) inadequate imaging contrast in the brainstem area, or c) inaccuracies stemming from the nonlinear spatial transformations applied to the imaging data.

### 3. Supplementary Table 1: The *task* model first-level specification and GLM contrasts performed in the analysis described in item 3.2.1 to 3.2.3

| Predictor | Reward Type | Presentation Frequency | Stimulus Type | One-sample *T*-test contrast coding | | | |
|---|---|---|---|---|---|---|---|
| | | | | **3.2.1.1.** Infreq. vs. freq. presented scenes | **3.2.1.2.** Infreq. vs. freq. presented feedbacks | **3.2.2.1.** Reward vs. neutral scenes | **3.2.2.2.** Reward vs. neutral feedbacks |
| 1 | Reward | Frequent | Scenes | -1 | | 1 | |
| 2 | Neutral | Infrequent | Scenes | 1 | | -1 | |
| 3 | Reward | Infrequent | Scenes | 1 | | 1 | |
| 4 | Neutral | Frequent | Scenes | -1 | | -1 | |
| 5 | Reward | Frequent | Feedbacks | | -1 | | 1 |
| 6 | Neutral | Infrequent | Feedbacks | | 1 | | -1 |
| 7 | Reward | Infrequent | Feedbacks | | 1 | | 1 |
| 8 | Neutral | Frequent | Feedbacks | | -1 | | -1 |
| 9 | - | - | Fixation cross | | | | |
| 10 | - | - | Button press | | | | |
| 11-16 | - | - | Realignment parameters | | | | |
| 17-22 | - | - | Physiological noise parameters | | | | |
| 23 | - | - | Intersession markers | | | | |
| 24 | - | - | Intercept | | | | |

**Supplementary Table 1. This table outlines the details of the '*task*' model specification and GLM one-sample *T*-test contrasts performed in the fMRI analysis described in item 3.2.1 to 3.2.2.** The predictor properties include Reward Type (Reward or Neutral), Presentation Frequency (Frequent or Infrequent), and Stimulus Type (Scenes or Feedbacks). Additionally, control predictors such as Fixation Cross, Button Press, Realignment Parameters (11-16), Physiological

Noise Parameters (17-22), Intersession Markers indicating concatenating of two fMRI sessions (23), and an Intercept (24) are incorporated to account for non-task-related brain activity and physiological artifacts. Any empty cells in this table were coded as 0 in the first-level analysis.

### 4. Supplementary Table 2: The *subsequent memory* model full-factorial ANOVA specification and GLM contrasts performed in the analysis described in item 3.2.3.3

| Cells | Reward Type | Presentation Frequency | Memory Outcome | Full factorial ANOVA *T*-test contrast coding | | | |
|---|---|---|---|---|---|---|---|
| | | | | Positive interaction effect between Frequency and Reward | Positive interaction effect between Reward and Memory Outcome | Positive interaction effect between Frequency and Memory Outcome | Positive interaction effect between Frequency, Reward, and Memory Outcome |
| 1 | Reward | Infrequent | Remembered | 1 | 1 | 1 | 1 |
| 2 | Reward | Infrequent | Forgotten | 1 | -1 | -1 | -1 |
| 3 | Neutral | Infrequent | Remembered | -1 | -1 | 1 | -1 |
| 4 | Neutral | Infrequent | Forgotten | -1 | 1 | -1 | 1 |
| 5 | Reward | Frequent | Remembered | -1 | 1 | -1 | -1 |
| 6 | Reward | Frequent | Forgotten | -1 | -1 | 1 | 1 |
| 7 | Neutral | Frequent | Remembered | 1 | -1 | -1 | 1 |
| 8 | Neutral | Frequent | Forgotten | 1 | 1 | 1 | -1 |

**Supplementary Table 2. This table outlines the details of the '*subsequent memory*' full factorial model specification and GLM contrasts performed in the fMRI analysis described in item 3.2.5.** In this full-factorial ANOVA, only scene stimulus presentation timepoint was inspected to refine the results to memory outcome only. Each predictor cell included properties of Reward Type (Reward or Neutral), Presentation Frequency (Frequent or Infrequent), and Memory Outcome in subsequent tests (Remembered or Forgotten). Additionally, in the first-level GLM utilised in this analysis, control predictors such as Fixation Cross, Button Press, Realignment Parameters (11-16), Physiological Noise Parameters (17-22), Intersession Markers indicating concatenating of two fMRI sessions (23), and an Intercept (24) are incorporated to account for non-task-related brain activity and physiological artifacts. Any empty cells in this table were coded as 0 in the first-level analysis.

### 5. Supplementary Table 3: The *subsequent memory* model first-level specification and GLM contrasts performed in the analysis described in item 3.2.3

| | | | | **One-sample *T*-test contrast coding** | | | |
|---|---|---|---|---|---|---|---|
| **Predictor** | **Reward Type** | **Presentation Frequency** | **Memory Outcome** | **3.2.3.1.** Remembered infreq. vs remembered freq. scenes | **3.2.3.2.** Remembered reward. Vs rememberd neutral scenes | **3.2.3.4.** Remembered infreq. vs. forgotten infreq. scenes | **3.2.3.5.** Remembered reward vs. forgotten reward scenes |
| 1 | Reward | Infrequent | Remembered | 1 | 1 | 1 | 1 |
| 2 | Reward | Frequent | Remembered | -1 | 1 | | 1 |
| 3 | Neutral | Infrequent | Remembered | 1 | -1 | 1 | |
| 4 | Neutral | Frequent | Remembered | -1 | -1 | | |
| 5 | Reward | Infrequent | Forgotten | | | -1 | -1 |
| 6 | Reward | Frequent | Forgotten | | | | -1 |
| 7 | Neutral | Infrequent | Forgotten | | | -1 | |
| 8 | Neutral | Frequent | Forgotten | | | | |
| 9 | - | - | Fixation cross | | | | |
| 10 | - | - | Button press | | | | |
| 11-16 | - | - | Realignment parameters | | | | |
| 17-22 | - | - | Physiological noise parameters | | | | |
| 23 | - | - | Intersession markers | | | | |
| 24 | - | - | Intercept | | | | |

**Supplementary Table 3. This table outlines the details of the '*subsequent memory*' model specification and GLM contrasts performed in the fMRI analysis described in item 3.2.3.** To isolate the effect of the two saliency types on encoding and to mitigate increased model complexity and the resultant decrease in statistical efficiency, only scene stimuli timepoints were analysed in first-level models. Each first-level model encompassed Reward Type (Reward or Neutral), Presentation Frequency (Frequent or Infrequent), and Memory Outcome in subsequent tests (Remembered or Forgotten). Additionally, control predictors such as Fixation Cross, Button Press, Realignment Parameters (11-16), Physiological Noise Parameters (17-22), Intersession Markers indicating concatenating of two fMRI sessions (23), and an Intercept (24) are incorporated to account for non-task-related brain activity and physiological artifacts. Any empty cells in this table were coded as 0 in the first-level analysis.

### 6. Supplementary Results 1: Interaction effects of delay length and two different salience types on false alarm (FA) and hit rates in recognition memory tests

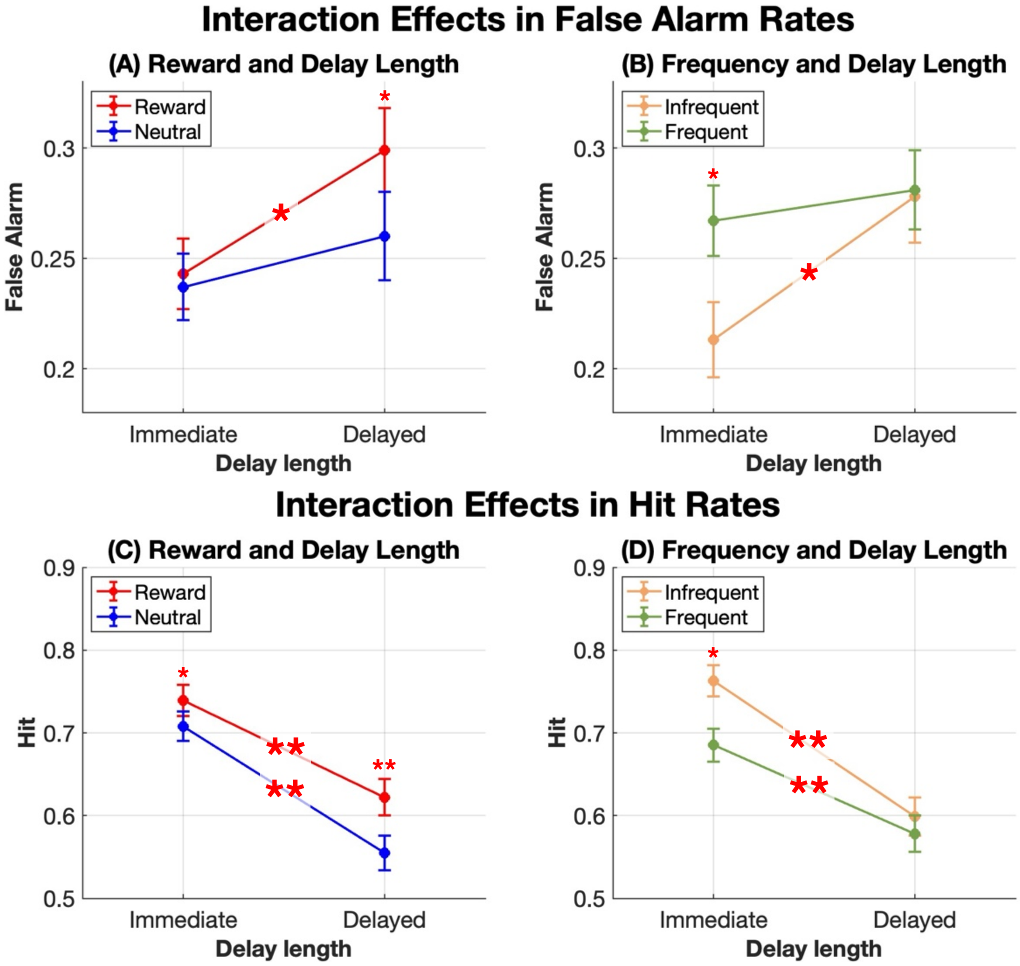


**Supplementary Figure 3. Profile plots of two-way interactions between delay length and both reward and contextual unexpectedness on FAs and hits. (A,C)** The interaction between delay length and reward-associated salience on FA **(A)** and hit **(C)** rates is shown. The red line represents the reward condition, while the blue line represents the neutral condition. **(B,D)** The interaction between delay length and frequency condition on FA **(B)** and hits **(D)** is shown. The green line represents the Frequent condition, while the orange line represents the Infrequent condition. For all panels, whiskers indicate standard error, and one asterisk (*) indicates significant differences with $p$<.05, and two asterisks (**) indicates significant differences with $p$<.001.

Significant two-way interactions were found between delay length and the two different salience types on FAs from a three-way repeated measures ANOVA. Reward-associated scenes show a more substantial increase in FAs over time compared to the neutral scenes (Supplementary Figure 3 A), $F(1,42)=4.137$, $p=.048$, $\eta_p^2=.090$, leading to significant difference in FA during delayed memory tests,

$F$(1,42)=4.699, $p$=.036, while infrequently presented scenes show lower FAs during immediate memory tests, $F$(1,42)=10.937, $p$=.002, but the difference diminishes over time (Supplementary Figure 3B), $F$(1,42)=6.995, $p$=.011, $\eta_p^2$=.143.

In the analysis of hit rate, from a three-way repeated measures ANOVA, the interaction between delay length and reward showed a trend towards significance (Supplementary Figure 3C), $F$(1,42)=3.711, $p$=.061, $\eta_p^2$=.081, indicating a more pronounced effect of reward on hit rate in reward-associated scenes in the delayed memory test, compared to the immediate test, $F$(1,42)=21.641, $p$<.001. In addition, the interaction between delay length and contextual unexpectedness was significant (Supplementary Figure 3D), $F$(1,42)=6.088, $p$=.018, $\eta_p^2$=.127, showing that the initial hit rate advantage from contextual unexpectedness in the immediate test ($F$[1,42]=10.937, $p$=.002) did not persist into the delayed test.

These results align with previous findings suggesting that associations with rewards predominantly influence decision biases rather than memory discrimination. Bowen et al. (4) demonstrated that while reward-associated stimuli can increase hit rates, they do not necessarily enhance memory discriminability (*D'*). This occurs because reward salience primarily affects decision-making tendencies, leading to a more liberal response bias towards reward-associated stimuli during recognition tests. Our results similarly show that participants exhibited better recognition of familiar reward-associated scenes (Supplementary Figure 4C and 4D), but this was counterbalanced by a larger increase in false alarms for these scenes (Supplementary Figure 4A and 4B). Consequently, there was no overall change in D'. This pattern suggests that reward motivation enhances both hit and false alarm rates, reinforcing the idea that it affects decision biases rather than memory discrimination.

### 7. Supplementary Results 2: Subsequently remembered trials vs. forgotten trials

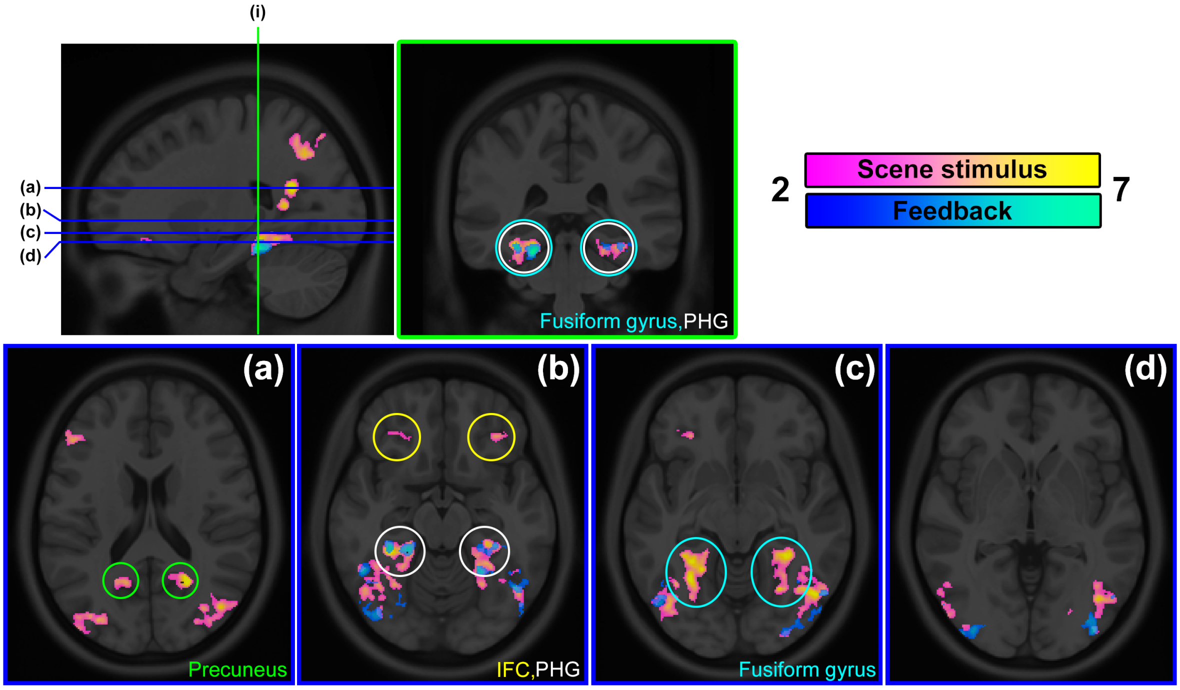


**Supplementary Figure 4. fMRI results from the subsequently remembered versus forgotten scenes.** All activations met a significance threshold of $p_{uncorr}<.001$ and were FDR-controlled. Axial slice (a) shows bilateral activation in the precuneus and the middle occipital gyrus during scene presentation. Axial slice (b) reveals bilateral inferior frontal cortex (IFC) activation during scene presentation and bilateral parahippocampal gyrus and inferior occipital lobe activations during both scene and feedback presentations. Axial slice (c) presents bilateral fusiform gyrus and ITL activations during scene presentation, and bilateral ITL activations during feedback presentation. Additionally, sagittal slice (c) displays left hippocampus activation during scene presentation. Finally, axial slice (d) demonstrates bilateral inferior occipital lobe activation during both scene and feedback presentations.

**During the *scene* presentation**, bilateral precuneus, PHG, middle and inferior occipital gyrus, IFC, fusiform gyrus, and the left hippocampus exhibited heightened activation for scenes that were later recognised, in contrast to those that were not, in both memory assessments (see Supplementary Figure 3). The midbrain and brainstem did not show notable activation.

These activation patterns suggest effective encoding and retrieval of visual stimuli, as evidenced by the studies that found the involvement of the precuneus, PHG, and middle occipital gyrus is linked to visuospatial processing, mental imagery, visual perception, attention, and memory consolidation (5,5–9). In addition, the roles of the hippocampus and IFC in episodic memory encoding and retrieval further corroborate the successful encoding evident in the subsequent memory test outcomes (8,10).

**During *feedback* presentation**, significant activations were observed in the bilateral occipital lobes in the middle and inferior areas, bilateral ITL, and left PHG. When using the midbrain and brainstem mask as an inclusive mask, no significant activation clusters were detected.

### 8. Supplementary Results 3: Interaction effects with reward and contextual unexpectedness in subsequently remembered scenes vs. forgotten scenes

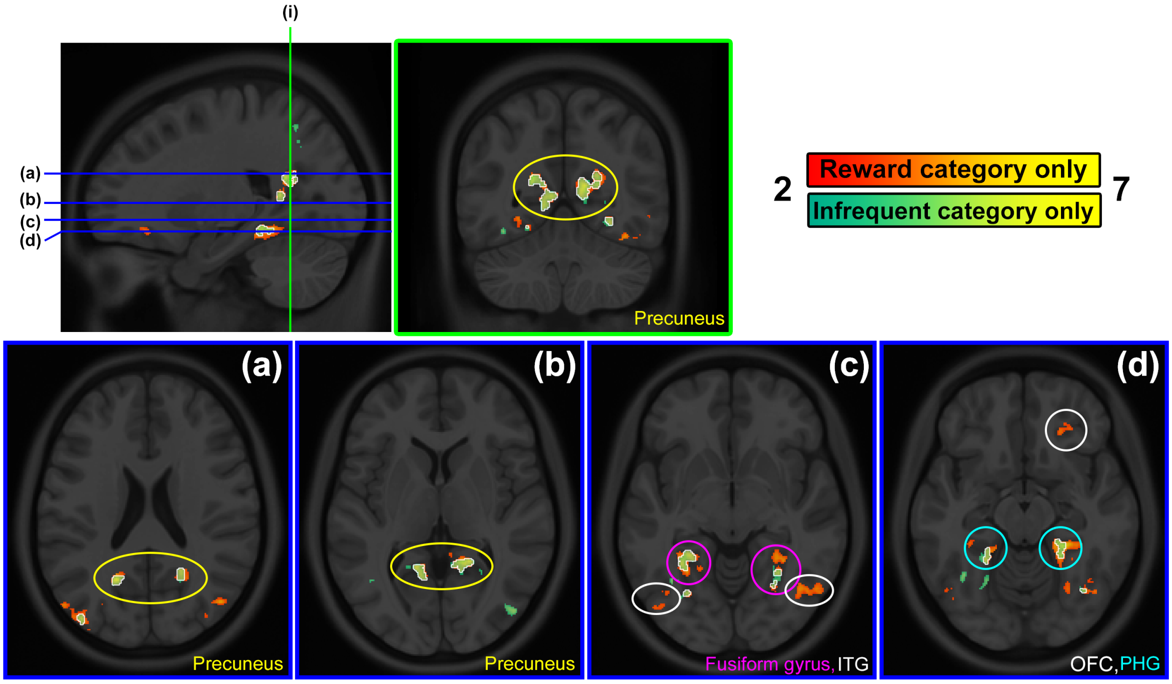


**Supplementary Figure 5. fMRI results from subsequently remembered versus forgotten scenes.** All activations were found with significance threshold of $p_{uncorr}$<.001 and was FDR-controlled. Activations encased with white lines delineate the overlapping voxels between the activations found from the two salience category contrasts. For activations specific **to infrequently presented scene**, axial slices (a), (b), and coronal slice (i) show activation in the bilateral precuneus. Slice (b) also shows bilateral middle occipital lobes alongside the bilateral precuneus. Slice (c) reveals bilateral fusiform gyrus and bilateral ITL, while slice (d) displays bilateral PHG activation. For **reward-associated scene**, activation patterns closely resemble those during infrequently presented scene, with the exception of right orbitofrontal cortex activation (OFC), as seen in axial slice (d). These overlapping activation patterns for remembered versus forgotten items in both salience categories—reward and contextual unexpectedness—address a potential critique concerning within-category competition. If memory encoding was affected by competition within the reward-associated category due to its predictability and more frequent presentation in some sessions, we might expect different activation patterns for successfully remembered items between the two categories. However, the observed similarity suggests that the type of salience, rather than internal category competition, is the primary driver of these memory-related activations.

#### 8.1. Subsequently remembered infrequent scenes vs. forgotten infrequent scenes

To examine memory effects in different salience types, trials with remembered versus forgotten scenes were contrasted separately for the two salient stimulus conditions: infrequent scenes and rewarded scenes. During ***infrequently*** presented scenes, significant activation differences between subsequently remembered and forgotten trials were observed in the bilateral calcarine sulcus, left fusiform gyrus, left lingual gyrus, right superior occipital lobe, and right fusiform gyrus (Supplementary Figure 5A).

**8.2. Subsequently remembered reward-associated scenes vs. forgotten reward-associated scenes**

A similar activation pattern was also identified for subsequently remembered ***reward-associated*** scenes, particularly in the right fusiform gyrus, left calcarine sulcus, and right superior occipital lobe, among other areas (see Supplementary Table 5 for more detailed results on both contrasts).

The activity patterns of successfully encoded scenes from both salience types show significant overlap (Supplementary Figure 5A; Jaccard Index = 0.5807, indicating that 58.07% of the activation in the smaller, contextually unexpected scenes' map is shared with the larger, reward-associated scenes' activation map). Despite this overlap, only the scenes that were contextually unexpected were associated with enhanced memory while rewarded scenes were not. This highlights that similar neural activity during encoding does not necessarily equate to uniform memory outcomes across different types of salience. Such a pattern implies that the better memory performance observed for contextually unexpected scenes may not solely come from differences in internal competition of each category. Rather, these shared activation profiles may represent common mechanisms of attention and memory that function across various types of stimuli. Further implications of these observations will be discussed in the subsequent Discussion section.

### 9. Supplementary Table 4-5: Lists of all fMRI activations

**Supplementary Table 4. fMRI activations, detailed statistics, and MNI coordinates of reward-associated and contextually unexpected category contrasts.** Equivk stands for equivalent cluster size, and equivZ stands for equivalent Z-score. FWE stands for family-wise error, and FDR stands for false discovery rate.

| | | Cluster | | | Peak | | | | MNI | | |
|---|---|---|---|---|---|---|---|---|---|---|---|
| **Side** | **Area** | ***p*(FWE-corr)** | ***p*(FDR-corr)** | **equivk** | ***p*(FWE-corr)** | ***p*(FDR-corr)** | **T** | **equivZ** | **x** | **y** | **z** |
| **Infrequently presented versus frequently presented *scene* stimuli** | | | | | | | | | | | |
| L | Insular cortex | 0 | 0 | 1424 | 0.057 | 0.052 | 6.47 | 5.43 | -37 | 23 | 0 |
| L | Fusiform gyrus | 0 | 0 | 3366 | 0.136 | 0.052 | 6.17 | 5.24 | -34 | -60 | -14 |
| R | Fusiform gyrus | 0 | 0 | 2225 | 0.147 | 0.052 | 6.14 | 5.22 | 38 | -46 | -22 |
| R | Inferior orbitofrontal cortex | 0.008 | 0.002 | 284 | 0.256 | 0.056 | 5.93 | 5.08 | 34 | 39 | -12 |
| R | Postcentral gyrus | 0.001 | 0 | 378 | 0.339 | 0.06 | 5.81 | 5.01 | 47 | -23 | 40 |
| R | Insular cortex | 0 | 0 | 1053 | 0.577 | 0.086 | 5.56 | 4.84 | 39 | 22 | -4 |
| L | Inferior parietal lobe & postcentral gyrus | 0 | 0 | 1186 | 0.592 | 0.086 | 5.55 | 4.83 | -52 | -27 | 57 |
| L | Medial superior frontal cortex | 0.044 | 0.002 | 215 | 0.643 | 0.086 | 5.5 | 4.8 | -3 | 36 | 43 |
| R | Caudate | 0.001 | 0 | 359 | 0.68 | 0.086 | 5.46 | 4.77 | 12 | 9 | 10 |
| L | Inferior parietal lobe | 0 | 0 | 732 | 0.729 | 0.089 | 5.42 | 4.74 | -45 | -45 | 52 |
| L | Inferior frontal cortex, pars opercularis | 0 | 0 | 622 | 0.907 | 0.138 | 5.21 | 4.59 | -47 | 5 | 31 |
| L | Superior frontal cortex | 0.771 | 0.043 | 100 | 0.995 | 0.207 | 4.91 | 4.38 | -25 | -8 | 48 |
| L | Inferior orbitofrontal cortex | 0.783 | 0.045 | 99 | 0.998 | 0.235 | 4.86 | 4.34 | -37 | 36 | -13 |
| R | Inferior parietal lobe | 0.143 | 0.018 | 171 | 0.999 | 0.249 | 4.83 | 4.32 | 35 | -50 | 52 |
| L | Inferior frontal cortex | 0.062 | 0.003 | 202 | 0.999 | 0.249 | 4.79 | 4.29 | -40 | 33 | 13 |
| L | Fusiform gyrus | 0.47 | 0.022 | 125 | 1 | 0.249 | 4.78 | 4.29 | -35 | -35 | -17 |
| L | Middle occipital lobe | 0.848 | 0.048 | 93 | 1 | 0.249 | 4.77 | 4.28 | -35 | -83 | 14 |
| R | Inferior frontal cortex, pars opercularis | 0 | 0 | 756 | 1 | 0.25 | 4.75 | 4.26 | 39 | 9 | 28 |
| L | Fusiform gyrus & parahippocampal gyrus | 0.033 | 0.002 | 226 | 1 | 0.258 | 4.73 | 4.25 | -23 | -35 | -19 |
| L | Calcarine sulcus | 0.686 | 0.036 | 107 | 1 | 0.27 | 4.7 | 4.23 | -24 | -59 | 10 |
| R | Inferior frontal cortex | 0.001 | 0 | 378 | 1 | 0.315 | 4.6 | 4.15 | 41 | 36 | 15 |

| | | | | | | | | | | | |
|---|---|---|---|---|---|---|---|---|---|---|---|
| R | Inferior occipital lobe & inferior temporal lobe | 0.003 | 0 | 324 | 1 | 0.315 | 4.58 | 4.13 | 46 | -79 | -3 |
| L | Caudate & pallidum | 0.053 | 0.003 | 208 | 1 | 0.315 | 4.56 | 4.12 | -8 | 12 | 5 |
| L | Superior parietal lobe | 0.006 | 0 | 292 | 1 | 0.34 | 4.45 | 4.03 | -26 | -71 | 59 |
| L | Inferior frontal cortex | 0.388 | 0.018 | 133 | 1 | 0.34 | 4.43 | 4.02 | -39 | 23 | 24 |
| R | Calcarine sulcus & precuneus | 0.04 | 0.002 | 219 | 1 | 0.42 | 4.2 | 3.84 | 16 | -53 | 10 |
| L | Middle occipital lobe | 0.481 | 0.029 | 124 | 1 | 0.435 | 4.18 | 3.82 | -30 | -81 | 27 |
| R | Anterior cingulate cortex (sup) | 0.817 | 0.045 | 96 | 1 | 0.538 | 3.99 | 3.68 | 11 | 26 | 27 |
| **Infrequently presented versus frequently presented *feedback*** | | | | | | | | | | | |
| L | Insular cortex | 0 | 0 | 1798 | 0 | 0 | 9.42 | 7 | -36 | 22 | -5 |
| R | Fusiform gyrus | 0 | 0 | 16219 | 0 | 0 | 8.42 | 6.52 | 47 | -60 | -15 |
| L | Inferior parietal lobe | 0 | 0 | 7190 | 0 | 0 | 8.05 | 6.33 | -31 | -54 | 39 |
| R | Inferior parietal lobe | 0 | 0 | 10037 | 0.001 | 0 | 7.7 | 6.14 | 34 | -61 | 46 |
| L | Fusiform gyrus | 0 | 0 | 16924 | 0.008 | 0.001 | 7.13 | 5.82 | -44 | -71 | -9 |
| R | Insular cortex | 0 | 0 | 2036 | 0.009 | 0.001 | 7.07 | 5.79 | 40 | 24 | -3 |
| R | Inferior frontal cortex, pars opercularis | 0 | 0 | 5802 | 0.011 | 0.001 | 7.01 | 5.75 | 43 | 35 | 19 |
| R | Caudate | 0 | 0 | 754 | 0.018 | 0.001 | 6.84 | 5.65 | 10 | 11 | 10 |
| L | Cerebellum (Crus 1) | 0 | 0 | 1187 | 0.048 | 0.003 | 6.52 | 5.46 | -10 | -73 | -25 |
| L&R | Posterior cingulate cortex | 0 | 0 | 466 | 0.05 | 0.003 | 6.5 | 5.45 | 2 | -34 | 26 |
| L | Caudate | 0 | 0 | 742 | 0.115 | 0.005 | 6.21 | 5.26 | -10 | 15 | 3 |
| L | Middle lateral prefrontal cortex | 0 | 0 | 847 | 0.208 | 0.007 | 5.99 | 5.13 | -48 | 52 | -9 |
| L | Inferior frontal cortex, pars opercularis | 0 | 0 | 925 | 0.474 | 0.015 | 5.65 | 4.9 | -45 | 23 | 22 |
| R | Inferior lateral orbitofrontal cortex | 0.006 | 0 | 306 | 0.707 | 0.023 | 5.42 | 4.74 | 41 | 43 | -7 |
| L | Middle temporal lobe | 0.002 | 0 | 347 | 0.997 | 0.076 | 4.86 | 4.34 | -62 | -41 | -7 |
| L | Medial superior frontal cortex | 0.002 | 0 | 354 | 0.998 | 0.078 | 4.84 | 4.33 | -4 | 40 | 49 |
| R | Medial superior frontal cortex | 0 | 0 | 493 | 0.998 | 0.079 | 4.84 | 4.33 | 2 | 39 | 44 |
| R | Parahippocampal gyrus | 0.856 | 0.05 | 95 | 1 | 0.095 | 4.73 | 4.25 | 33 | -24 | -22 |
| R | Precuneus | 0 | 0 | 545 | 1 | 0.098 | 4.71 | 4.23 | 8 | -75 | 51 |
| L | Medial orbitofrontal cortex | 0.043 | 0.005 | 223 | 1 | 0.11 | 4.65 | 4.19 | -15 | 59 | -20 |
| L | Superior frontal cortex | 0.013 | 0 | 272 | 1 | 0.15 | 4.47 | 4.06 | 21 | 64 | -13 |

| | | | | | | | | | | | |
|---|---|---|---|---|---|---|---|---|---|---|---|
| L | Precuneus | 0.503 | 0.019 | 126 | 1 | 0.159 | 4.44 | 4.03 | -8 | -72 | 45 |
| R | Superior temporal lobe | 0.737 | 0.036 | 106 | 1 | 0.172 | 4.4 | 4 | 47 | -26 | -4 |
| **Reward-associated versus neutral *scene* stimuli** | | | | | | | | | | | |
| L | Superior parietal lobe | 0.1 | 0.039 | 186 | 1 | 1 | 4.51 | 4.09 | -18 | -64 | 55 |
| **Reward-associated versus neutral *feedback*** | | | | | | | | | | | |
| L | Inferior & middle occipital lobe & fusiform gyrus | 0 | 0 | 21985 | 0 | 0 | 10.15 | 7.32 | -38 | -90 | -6 |
| R | Middle occipital lobe | 0 | 0 | 52255 | 0 | 0 | 9.71 | 7.13 | 33 | -68 | 28 |
| R | Inferior frontal cortex, pars opercularis | 0 | 0 | 3732 | 0.001 | 0 | 7.64 | 6.11 | 46 | 9 | 26 |
| L | Inferior & superior parietal lobe | 0 | 0 | 7675 | 0.003 | 0 | 7.39 | 5.97 | -26 | -63 | 36 |
| L&R | Middle cingulate cortex | 0 | 0 | 1214 | 0.023 | 0 | 6.76 | 5.61 | 4 | -4 | 30 |
| R | Ventral striatum | 0 | 0 | 1483 | 0.036 | 0.001 | 6.6 | 5.51 | 7 | 8 | -9 |
| R | Insular cortex | 0.117 | 0.004 | 188 | 0.065 | 0.001 | 6.4 | 5.39 | 34 | 17 | 1 |
| R | Superior frontal cortex | 0 | 0 | 609 | 0.18 | 0.003 | 6.04 | 5.16 | 26 | 0 | 53 |
| L | Ventral striatum | 0 | 0 | 1093 | 0.319 | 0.004 | 5.82 | 5.01 | -11 | 14 | -5 |
| R | Anterior orbitofrontal cortex | 0 | 0 | 501 | 0.916 | 0.022 | 5.17 | 4.57 | 23 | 42 | -15 |
| R | Anterior cingulate cortex (pre) | 0.359 | 0.013 | 143 | 0.974 | 0.032 | 5.03 | 4.47 | 13 | 40 | 20 |
| L | Anterior cingulate cortex (pre) | 0.376 | 0.035 | 141 | 0.987 | 0.037 | 4.97 | 4.42 | -4 | 38 | -1 |
| R | Medial superior frontal cortex | 0.07 | 0.004 | 208 | 1 | 0.066 | 4.73 | 4.25 | 3 | 22 | 44 |
| R | Inferior orbitofrontal cortex | 0.12 | 0.004 | 187 | 1 | 0.081 | 4.65 | 4.19 | 32 | 29 | -9 |
| L | Inferior frontal cortex, pars opercularis | 0.038 | 0.002 | 232 | 1 | 0.097 | 4.57 | 4.13 | -47 | 8 | 26 |
| R | Superior frontal cortex | 0.778 | 0.039 | 104 | 1 | 0.102 | 4.55 | 4.11 | 25 | 66 | 1 |
| L | Inferior temporal lobe | 0.275 | 0.01 | 154 | 1 | 0.138 | 4.4 | 4 | -48 | -47 | -15 |
| R | Middle frontal lobe | 0 | 0 | 619 | 1 | 0.168 | 4.3 | 3.92 | 40 | 39 | 15 |
| L&R | Anterior cingulate cortex (pre & sup) | 0.334 | 0.013 | 146 | 1 | 0.181 | 4.26 | 3.89 | -1 | 41 | 9 |
| R | Medial orbitofrontal cortex & anterior cingulate cortex (pre) | 0.123 | 0.007 | 186 | 1 | 0.343 | 3.93 | 3.63 | 3 | 46 | -4 |
| **Infrequently presented reward-associated versus frequently presented reward-associated *scene* stimuli** | | | | | | | | | | | |
| L | Insular cortex | 0 | 0.001 | 674 | 0.093 | 0.084 | 6.3 | 5.32 | -30 | 30 | 2 |
| L | Calcarine sulcus | 0.107 | 0.02 | 184 | 0.497 | 0.225 | 5.63 | 4.89 | -19 | -62 | 7 |

| | | | | | | | | | | | |
|---|---|---|---|---|---|---|---|---|---|---|---|
| L | Superior parietal lobe | 0.125 | 0.048 | 178 | 0.542 | 0.225 | 5.59 | 4.86 | -22 | -51 | 61 |
| L | Precentral gyrus | 0.339 | 0.048 | 140 | 0.987 | 0.754 | 4.98 | 4.43 | -29 | -6 | 47 |
| L | Postcentral gyrus | 0.074 | 0.02 | 198 | 1 | 0.802 | 4.71 | 4.23 | -52 | -24 | 58 |
| R | Insular cortex | 0.322 | 0.048 | 142 | 1 | 0.802 | 4.26 | 3.89 | 30 | 26 | -7 |
| **Infrequently presented reward versus frequently presented reward *feedback*** | | | | | | | | | | | |
| R | Angular gyrus | 0 | 0 | 3733 | 0.01 | 0.012 | 7.04 | 5.77 | 30 | -56 | 43 |
| L | Inferior parietal lobe | 0 | 0 | 2547 | 0.043 | 0.017 | 6.55 | 5.48 | -42 | -51 | 50 |
| R | Insular cortex | 0 | 0 | 611 | 0.094 | 0.026 | 6.28 | 5.31 | 30 | 27 | -4 |
| L | Inferior occipital lobe | 0 | 0 | 3762 | 0.111 | 0.026 | 6.23 | 5.27 | -45 | -64 | -13 |
| R | Inferior temporal lobe | 0 | 0 | 3201 | 0.16 | 0.034 | 6.09 | 5.19 | 56 | -55 | -11 |
| L | Inferior parietal lobe | 0 | 0 | 600 | 0.205 | 0.04 | 6 | 5.13 | -56 | -33 | 51 |
| L | Insular cortex | 0 | 0 | 459 | 0.828 | 0.148 | 5.3 | 4.66 | -32 | 20 | 4 |
| R | Inferior frontal cortex, pars opercularis | 0 | 0 | 1454 | 0.958 | 0.208 | 5.09 | 4.51 | 58 | 16 | 22 |
| L&R | Posterior & middle cingulate cortex | 0.107 | 0.01 | 187 | 0.992 | 0.237 | 4.94 | 4.4 | 4 | -35 | 25 |
| R | Supramarginal | 0.646 | 0.035 | 113 | 0.999 | 0.255 | 4.81 | 4.31 | 44 | -36 | 44 |
| R | Medial superior frontal cortex | 0.316 | 0.017 | 145 | 1 | 0.311 | 4.71 | 4.23 | 2 | 32 | 50 |
| R | Inferior occipital lobe | 0.001 | 0 | 405 | 1 | 0.311 | 4.61 | 4.16 | 40 | -85 | -3 |
| R | Inferior parietal lobe | 0.049 | 0.005 | 217 | 1 | 0.311 | 4.61 | 4.15 | 51 | -44 | 55 |
| L | Cerebellum (Crus 2) | 0.349 | 0.017 | 141 | 1 | 0.311 | 4.6 | 4.15 | -8 | -77 | -31 |
| R | Fusiform gyrus | 0.002 | 0.002 | 359 | 1 | 0.311 | 4.59 | 4.14 | 35 | -38 | -22 |
| L | Fusiform gyrus | 0.001 | 0.002 | 375 | 1 | 0.34 | 4.54 | 4.1 | -36 | -50 | -24 |
| R | Middle lateral prefrontal cortex | 0.598 | 0.032 | 117 | 1 | 0.394 | 4.43 | 4.02 | 41 | 46 | 17 |
| R | Inferior frontal cortex, pars opercularis | 0 | 0 | 555 | 1 | 0.396 | 4.37 | 3.98 | 42 | 23 | 22 |
| R | Cerebellum VI | 0.357 | 0.025 | 140 | 1 | 0.434 | 4.32 | 3.94 | 35 | -78 | -21 |
| R | Middle lateral prefrontal cortex | 0.286 | 0.014 | 149 | 1 | 0.434 | 4.29 | 3.91 | 45 | 49 | 1 |
| R | Inferior parietal lobe | 0.205 | 0.01 | 162 | 1 | 0.502 | 4.13 | 3.79 | 48 | -49 | 45 |
| L | Middle occipital lobe | 0.54 | 0.032 | 122 | 1 | 0.544 | 3.93 | 3.63 | -29 | -71 | 30 |

**Supplementary Table 5. fMRI activations, detailed statistics, and MNI coordinates of subsequent-memory-associated contrasts.** Equivk stands for equivalent cluster size, and equivZ stands for equivalent Z-score. FWE stands for family-wise error, and FDR stands for false discovery rate.

| | | Cluster | | | Peak | | | | MNI | | |
|---|---|---|---|---|---|---|---|---|---|---|---|
| **Side** | **Area** | ***p*(FWE-corr)** | ***p*(FDR-corr)** | **equivk** | ***p*(FWE-corr)** | ***p*(FDR-corr)** | **T** | **equivZ** | **x** | **y** | **z** |
| **Subsequently remembered versus forgotten *scene* stimuli** | | | | | | | | | | | |
| R | Fusiform gyrus, inferior temporal lobe, & parahippocampal gyrus | 0 | 0 | 9575 | 0 | 0 | 9.28 | 6.93 | 33 | -37 | -14 |
| R | Calcarine sulcus & Precuneus | 0 | 0 | 2472 | 0 | 0 | 9.28 | 6.93 | 24 | -57 | 20 |
| R | Middle occipital lobe | 0 | 0 | 7950 | 0 | 0 | 8.93 | 6.77 | 39 | -84 | 26 |
| L | Inferior frontal cortex | 0 | 0 | 998 | 0 | 0 | 8.07 | 6.34 | -47 | 33 | 15 |
| L | Fusiform gyrus | 0 | 0 | 8944 | 0.002 | 0.001 | 7.59 | 6.08 | -29 | -56 | -10 |
| L | Middle occipital lobe | 0 | 0 | 5141 | 0.004 | 0.001 | 7.32 | 5.93 | -23 | -69 | 26 |
| L | Hippocampus & parahippocampal gyrus | 0.228 | 0.013 | 152 | 0.248 | 0.01 | 5.94 | 5.09 | -21 | -16 | -22 |
| L | Calcarine sulcus & precuneus | 0 | 0 | 1655 | 0.262 | 0.011 | 5.92 | 5.08 | -11 | -51 | 3 |
| L | Cerebellum VI | 0.029 | 0.002 | 228 | 0.756 | 0.029 | 5.39 | 4.72 | -8 | -73 | -23 |
| L | Inferior frontal cortex, pars opercularis | 0.001 | 0 | 376 | 0.826 | 0.034 | 5.32 | 4.67 | -34 | 8 | 29 |
| L | Superior parietal lobe | 0 | 0 | 463 | 0.957 | 0.054 | 5.11 | 4.52 | -21 | -61 | 56 |
| R | Inferior parietal lobe | 0.031 | 0.002 | 226 | 0.981 | 0.063 | 5.03 | 4.47 | 29 | -51 | 46 |
| L | Inferior orbitofrontal cortex, anterior orbitofrontal cortex, & posterior orbitofrontal cortex | 0.001 | 0.002 | 349 | 0.981 | 0.063 | 5.03 | 4.46 | -34 | 34 | -15 |
| R | Inferior frontal cortex | 0.05 | 0.012 | 208 | 1 | 0.15 | 4.57 | 4.13 | 49 | 22 | 23 |
| R | Inferior occipital lobe | 0.199 | 0.012 | 157 | 1 | 0.297 | 4.2 | 3.85 | 32 | -87 | -6 |
| R | Anterior orbitofrontal cortex | 0.321 | 0.018 | 139 | 1 | 0.377 | 4.06 | 3.73 | 25 | 38 | -13 |
| **Subsequently remembered versus forgotten *feedback*** | | | | | | | | | | | |
| R | Fusiform gyrus & parahippocampal gyrus | 0 | 0 | 1512 | 0 | 0.001 | 8.03 | 6.31 | 23 | -36 | -19 |
| R | Middle & inferior occipital lobe & fusiform gyrus | 0 | 0 | 3223 | 0.003 | 0.002 | 7.39 | 5.97 | 41 | -72 | -18 |
| R | Inferior temporal lobe | 0 | 0 | 502 | 0.121 | 0.041 | 6.21 | 5.26 | 50 | -51 | -18 |
| L | Fusiform gyrus | 0 | 0 | 597 | 0.238 | 0.07 | 5.96 | 5.1 | -32 | -52 | -18 |
| L | Middle occipital lobe | 0 | 0 | 437 | 0.671 | 0.204 | 5.47 | 4.78 | -37 | -80 | 0 |

| | | | | | | | | | | | |
|---|---|---|---|---|---|---|---|---|---|---|---|
| R | Inferior temporal lobe | 0 | 0 | 919 | 0.956 | 0.299 | 5.11 | 4.52 | -53 | -61 | -20 |
| R | Middle occipital lobe | 0.001 | 0 | 393 | 0.964 | 0.299 | 5.09 | 4.51 | 33 | -87 | 28 |
| R | Cerebellum VI | 0.01 | 0.006 | 272 | 0.977 | 0.304 | 5.04 | 4.47 | 32 | -82 | -19 |
| R | Middle occipital lobe | 0.001 | 0 | 399 | 0.996 | 0.324 | 4.91 | 4.38 | 36 | -72 | 32 |
| L | Inferior occipital lobe & inferior temporal lobe | 0 | 0 | 631 | 0.996 | 0.324 | 4.9 | 4.38 | -53 | -68 | -6 |
| L | Fusiform gyrus & parahippocampal gyrus | 0 | 0 | 639 | 0.998 | 0.324 | 4.85 | 4.33 | -28 | -31 | -23 |
| L | Fusiform gyrus | 0.131 | 0.01 | 174 | 1 | 0.446 | 4.3 | 3.92 | -38 | -46 | -25 |
| **Subsequently remembered infrequently versus frequently presented *scene* stimuli** | | | | | | | | | | | |
| L | Calcarine sulcus & precuneus | 0.002 | 0.003 | 333 | 0.999 | 0.757 | 4.84 | 4.33 | -11 | -61 | 9 |
| L | Inferior parietal lobe & postcentral gyrus | 0.136 | 0.021 | 168 | 0.999 | 0.757 | 4.84 | 4.33 | -53 | -23 | 42 |
| R | Postcentral gyrus | 0.01 | 0.003 | 266 | 0.999 | 0.757 | 4.82 | 4.31 | 36 | -29 | 41 |
| R | Inferior frontal cortex, pars opercularis | 0.021 | 0.004 | 236 | 1 | 0.775 | 4.59 | 4.14 | 41 | 8 | 26 |
| L | Fusiform gyrus | 0.007 | 0.003 | 277 | 1 | 0.803 | 4.37 | 3.98 | -32 | -63 | -10 |
| L | Middle superior frontal cortex | 0.14 | 0.021 | 167 | 1 | 0.803 | 4.33 | 3.95 | -7 | 31 | 39 |
| **Subsequently remembered reward-associated versus neutral *scene* stimuli** | | | | | | | | | | | |
| L | Medial orbitofrontal cortex | 0.002 | 0.007 | 334 | 0.989 | 0.457 | 4.98 | 4.43 | -15 | 47 | -10 |
| **Subsequently remembered infrequently versus frequently presented *feedback*** | | | | | | | | | | | |
| L | Inferior parietal lobe & postcentral gyrus | 0 | 0 | 6937 | 0 | 0 | 8.42 | 6.52 | -34 | -55 | 42 |
| R | Fusiform gyrus, & inferior temporal lobe | 0 | 0 | 10265 | 0.001 | 0.001 | 7.94 | 6.27 | 47 | -53 | -14 |
| L | Fusiform gyrus & inferior occipital lobe | 0 | 0 | 11391 | 0.003 | 0.001 | 7.46 | 6.01 | -34 | -53 | -16 |
| R | Inferior parietal lobe | 0 | 0 | 8501 | 0.003 | 0.001 | 7.45 | 6 | 33 | -46 | 37 |
| L | Insular cortex | 0 | 0 | 1312 | 0.011 | 0.002 | 7.02 | 5.76 | -35 | 22 | -5 |
| R | Caudate | 0 | 0.001 | 508 | 0.021 | 0.003 | 6.81 | 5.63 | 10 | 11 | 10 |
| L&R | Posterior cingulate cortex | 0 | 0 | 662 | 0.028 | 0.003 | 6.71 | 5.57 | 1 | -34 | 26 |
| R | Insular cortex | 0 | 0 | 1621 | 0.028 | 0.003 | 6.7 | 5.57 | 29 | 25 | -6 |
| R | Inferior frontal cortex | 0 | 0 | 4949 | 0.046 | 0.004 | 6.54 | 5.47 | 44 | 38 | 13 |
| L | Middle superior frontal cortex | 0 | 0 | 1108 | 0.142 | 0.006 | 6.15 | 5.22 | -48 | 51 | -6 |
| R | Middle superior frontal cortex | 0 | 0 | 532 | 0.326 | 0.012 | 5.83 | 5.02 | 2 | 38 | 46 |
| L | Inferior frontal cortex | 0 | 0 | 646 | 0.602 | 0.023 | 5.54 | 4.82 | -49 | 25 | 24 |

| | | | | | | | | | | | |
|---|---|---|---|---|---|---|---|---|---|---|---|
| R | Middle temporal lobe | 0 | 0 | 809 | 0.693 | 0.028 | 5.45 | 4.76 | 66 | -33 | -9 |
| L | Middle superior frontal cortex | 0.011 | 0.001 | 272 | 0.926 | 0.048 | 5.17 | 4.57 | -2 | 39 | 47 |
| L | Cerebellum (Crus I) | 0 | 0 | 686 | 0.994 | 0.071 | 4.93 | 4.4 | -7 | -74 | -25 |
| R | Superior frontal cortex | 0.028 | 0.001 | 233 | 0.998 | 0.08 | 4.87 | 4.35 | 22 | 56 | -8 |
| L | Precentral gyrus | 0.044 | 0.002 | 216 | 1 | 0.159 | 4.5 | 4.08 | -43 | 8 | 28 |
| L | Middle temporal lobe | 0.192 | 0.006 | 161 | 1 | 0.182 | 4.43 | 4.02 | -63 | -40 | -6 |
| L | Caudate | 0.056 | 0.002 | 207 | 1 | 0.188 | 4.41 | 4.01 | -9 | 13 | 2 |
| R | Inferior orbitofrontal cortex | 0.198 | 0.017 | 160 | 1 | 0.24 | 4.28 | 3.9 | 45 | 49 | -11 |
| R | Superior temporal lobe | 0.467 | 0.017 | 126 | 1 | 0.24 | 4.27 | 3.9 | 48 | -26 | -5 |
| R | Precuenus | 0.068 | 0.002 | 200 | 1 | 0.249 | 4.25 | 3.89 | 8 | -72 | 45 |
| **Subsequently remembered reward-associated versus neutral *feedback*** | | | | | | | | | | | |
| R | Middle occipital lobe | 0 | 0 | 43522 | 0 | 0 | 9.54 | 7.05 | 29 | -64 | 35 |
| L | Inferior occipital lobe | 0 | 0 | 19437 | 0 | 0 | 9.2 | 6.89 | -37 | -91 | -6 |
| R | Inferior frontal cortex, pars opercularis | 0 | 0 | 3031 | 0.013 | 0.001 | 6.95 | 5.72 | 46 | 7 | 24 |
| L | Superior parietal lobe | 0 | 0 | 6147 | 0.018 | 0.001 | 6.84 | 5.65 | -27 | -67 | 30 |
| L | Anterior cinguate cortex (pre) | 0 | 0 | 2142 | 0.042 | 0.001 | 6.56 | 5.48 | -2 | 42 | 9 |
| R | Ventral striatum | 0 | 0 | 1998 | 0.048 | 0.001 | 6.52 | 5.46 | 8 | 9 | -11 |
| R | Insular cortex | 0.47 | 0.028 | 129 | 0.076 | 0.002 | 6.36 | 5.36 | 33 | 17 | 1 |
| R | Inferior orbitofrontal cortex | 0.005 | 0 | 308 | 0.295 | 0.006 | 5.86 | 5.04 | 29 | 22 | -10 |
| R | Anterior cingulate cortex (sup) | 0.067 | 0.006 | 206 | 0.342 | 0.007 | 5.8 | 5 | 4 | 14 | 23 |
| L&R | Middle cingulate cortex | 0 | 0 | 637 | 0.562 | 0.012 | 5.56 | 4.84 | 7 | -13 | 29 |
| L | Anterior cingulate cortex (sup) | 0.593 | 0.045 | 118 | 0.718 | 0.017 | 5.41 | 4.74 | -3 | 31 | 19 |
| R | Anterior orbitofrontal cortex | 0.116 | 0.006 | 185 | 0.726 | 0.018 | 5.41 | 4.73 | 24 | 45 | -14 |
| L | Ventral striatum | 0 | 0 | 1408 | 0.797 | 0.021 | 5.33 | 4.68 | -13 | 5 | -14 |
| L | Inferior frontal cortex, pars opercularis | 0.004 | 0 | 323 | 0.989 | 0.05 | 4.96 | 4.42 | -37 | 2 | 29 |
| R | Middle superior frontal cortex | 0.008 | 0 | 293 | 1 | 0.123 | 4.53 | 4.1 | 43 | 45 | 16 |
| R | Middle superior frontal cortex | 0.116 | 0.006 | 185 | 1 | 0.124 | 4.52 | 4.09 | 34 | -3 | 51 |
| R | Insular cortex | 0.41 | 0.028 | 135 | 1 | 0.139 | 4.47 | 4.05 | 44 | 18 | -9 |
| R | Middle superior frontal cortex | 0.513 | 0.03 | 125 | 1 | 0.18 | 4.35 | 3.96 | 45 | 37 | 23 |

| | | | | | | | | | | | |
|---|---|---|---|---|---|---|---|---|---|---|---|
| **Subsequently remembered versus forgotten infrequently presented *scene* stimuli** | | | | | | | | | | | |
| L | Calcarine sulcus & precuneus | 0 | 0 | 1406 | 0.492 | 0.239 | 5.67 | 4.9 | -8 | -60 | 8 |
| R | Calcarine sulcus & precuneus | 0 | 0 | 1114 | 0.799 | 0.239 | 5.37 | 4.69 | 13 | -56 | 15 |
| L | Fusiform gyrus & middle temporal lobe | 0.011 | 0.009 | 266 | 0.827 | 0.239 | 5.33 | 4.67 | -38 | -60 | -9 |
| R | Superior occipital lobe | 0 | 0 | 621 | 0.912 | 0.239 | 5.22 | 4.59 | 24 | -60 | 49 |
| L | Fusiform gyrus & lingual gyrus | 0 | 0 | 659 | 0.916 | 0.239 | 5.21 | 4.58 | -27 | -44 | -9 |
| R | Middle occipital lobe | 0.232 | 0.013 | 152 | 0.951 | 0.248 | 5.14 | 4.53 | 46 | -78 | 3 |
| R | Fusiform gyrus & parahippocampal gyrus | 0.135 | 0.009 | 172 | 0.971 | 0.26 | 5.08 | 4.49 | 24 | -39 | -11 |
| L | Lingual gyrus & fusiform gyrus | 0.003 | 0.001 | 324 | 0.994 | 0.297 | 4.95 | 4.4 | -27 | -64 | -11 |
| L | Middle occipital lobe | 0.011 | 0.001 | 268 | 0.997 | 0.297 | 4.9 | 4.36 | -42 | -84 | 23 |
| R | Fusiform gyrus & inferior temporal lobe | 0.001 | 0 | 365 | 1 | 0.297 | 4.77 | 4.26 | 38 | -41 | -21 |
| R | Fusiform gyrus & lingual gyrus | 0.011 | 0.001 | 267 | 1 | 0.329 | 4.68 | 4.2 | 29 | -59 | -8 |
| **Subsequently remembered versus forgotten reward-associated *scene* stimuli** | | | | | | | | | | | |
| R | Fusiform gyrus & parahippocampal gyrus | 0 | 0 | 4047 | 0.003 | 0.004 | 7.5 | 6.01 | 25 | -38 | -12 |
| L | Middle occipital lobe | 0 | 0 | 1499 | 0.009 | 0.006 | 7.12 | 5.79 | -41 | -84 | 25 |
| L | Calcarine sulcus & precuneus | 0 | 0 | 1248 | 0.019 | 0.008 | 6.88 | 5.66 | -20 | -57 | 20 |
| L | Fusiform gyrus & lingual gyrus | 0 | 0 | 1639 | 0.436 | 0.086 | 5.72 | 4.93 | -31 | -46 | -6 |
| R | Calcarin sulcus & precuneus | 0 | 0 | 1708 | 0.759 | 0.129 | 5.4 | 4.72 | 23 | -53 | 19 |
| R | Middle occipital lobe & middle temporal lobe | 0 | 0 | 536 | 0.955 | 0.183 | 5.12 | 4.52 | 46 | -73 | 22 |
| L | Middle temporal lobe & inferior occipital lobe | 0 | 0 | 414 | 1 | 0.27 | 4.78 | 4.28 | -53 | -70 | -2 |
| R | Inferior temporal lobe & fusiform gyrus | 0.001 | 0 | 358 | 1 | 0.27 | 4.76 | 4.26 | 44 | -52 | -18 |
| R | Superior occipital lobe | 0.002 | 0 | 341 | 1 | 0.282 | 4.67 | 4.19 | 27 | -69 | 38 |
| R | Anterior and posterior orbitofrontal cortex | 0.37 | 0.023 | 136 | 1 | 0.374 | 4.38 | 3.98 | 25 | 38 | -13 |
| R | Middle occipital lobe | 0.252 | 0.016 | 151 | 1 | 0.375 | 4.36 | 3.96 | 37 | -83 | 26 |
| L | Inferior occipital lobe | 0.007 | 0 | 289 | 1 | 0.462 | 4.11 | 3.77 | -45 | -72 | -13 |
| **Subsequently remembered versus forgotten infrequently presented *feedbacks*** | | | | | | | | | | | |
| R | Inferior frontal gyrus | 0.204 | 0.037 | 154 | 0.889 | 0.668 | 5.26 | 4.62 | 43 | 30 | 24 |
| L | Fusiform gyrus | 0.204 | 0.037 | 154 | 0.994 | 0.771 | 4.95 | 4.4 | -40 | -52 | -17 |
| R | Inferior occipital lobe | 0.323 | 0.042 | 137 | 0.998 | 0.771 | 4.88 | 4.35 | 47 | -77 | -15 |

| | | | | | | | | | | | |
|---|---|---|---|---|---|---|---|---|---|---|---|
| L | Inferior occipital lobe | 0.012 | 0.008 | 258 | 1 | 0.956 | 4.57 | 4.12 | -51 | -58 | -16 |
| R | Inferior temporal lobe | 0.025 | 0.008 | 230 | 1 | 0.956 | 4.52 | 4.08 | 54 | -64 | -17 |
| **Subsequently remembered versus forgotten reward *feedbacks*** | | | | | | | | | | | |
| *No suprathreshold clusters* | | | | | | | | | | | |

### References


1. Yi YJ, Lüsebrink F, Ludwig M, Maaß A, Ziegler G, Yakupov R, et al. It is the locus coeruleus! Or… is it?: a proposition for analyses and reporting standards for structural and functional magnetic resonance imaging of the noradrenergic locus coeruleus. Neurobiol Aging. 2023 Sep;129:137–48.

2. Fonov V, Evans AC, Botteron K, Almli CR, McKinstry RC, Collins DL. Unbiased average age-appropriate atlases for pediatric studies. NeuroImage. 2011 Jan;54(1):313–27.

3. Avants BB, Tustison NJ, Song G, Cook PA, Klein A, Gee JC. A reproducible evaluation of ANTs similarity metric performance in brain image registration. NeuroImage. 2011 Feb;54(3):2033–44.

4. Bowen HJ, Marchesi ML, Kensinger EA. Reward motivation influences response bias on a recognition memory task. Cognition. 2020 Oct;203:104337.

5. Cabeza R, Ciaramelli E, Olson IR, Moscovitch M. The parietal cortex and episodic memory: an attentional account. Nat Rev Neurosci. 2008 Aug;9(8):613–25.

6. Cavanna AE, Trimble MR. The precuneus: a review of its functional anatomy and behavioural correlates. Brain. 2006 Mar 1;129(3):564–83.

7. Lundstrom BN, Ingvar M, Petersson KM. The role of precuneus and left inferior frontal cortex during source memory episodic retrieval. NeuroImage. 2005 Oct;27(4):824–34.

8. Vilberg KL, Rugg MD. The Neural Correlates of Recollection: Transient Versus Sustained fMRI Effects. J Neurosci. 2012 Nov 7;32(45):15679–87.

9. Wagner AD, Schacter DL, Rotte M, Koutstaal W, Maril A, Dale AM, et al. Building Memories: Remembering and Forgetting of Verbal Experiences as Predicted by Brain Activity. Science. 1998 Aug 21;281(5380):1188–91.

10. Blumenfeld RS, Ranganath C. Prefrontal Cortex and Long-Term Memory Encoding: An Integrative Review of Findings from Neuropsychology and Neuroimaging. The Neuroscientist. 2007 Jun;13(3):280–91.
